# Contrasting effects of glutamate and branched-chain amino acid metabolism on acid tolerance in a *Castellaniella* isolate from acidic groundwater

**DOI:** 10.1101/2025.09.26.678905

**Authors:** Jennifer L. Goff, Konnor L. Durrence, Michael P. Thorgersen, Valentine V. Trotter, Yan Chen, Suzanne M. Kosina, Audrey L.W. Wang, Farris L. Poole, Trent R. Northen, Christopher J. Petzold, Adam M. Deutschbauer, Michael W. W. Adams

## Abstract

Groundwater acidification co-occurring with nitrate pollution is a common, global environmental health hazard. Denitrifying bacteria have been leveraged for the *in-situ* removal of nitrate in groundwater. However, co-existing stressors—like low pH—reduce the efficacy of biological removal processes. *Castellaniella* sp. str. MT123 is a complete denitrifier that was isolated from acidic, nitrate-contaminated groundwater. The strain grows robustly by nitrate respiration at pH < 6.0 completely reducing nitrate to dinitrogen gas. Genomic analyses of MT123 revealed few previously characterized acid tolerance genes. Thus, we utilized a combination of proteomics, metabolomics, and competitive mutant fitness to characterize the genetic mechanisms of MT123 acclimation to growth under mildly acidic conditions. We found that glutamate accumulation is critical in the acid acclimation of MT123, likely through consumption of intracellular protons via glutamate decarboxylation to GABA. This is despite the fact that MT123 lacks the canonical glutamate decarboxylase-glutamate/GABA antiporter system implicated in acid tolerance in other bacteria. Additionally, branched chain amino acid (BCAA) accumulation was detrimental to cell growth at lower pHs, possibly through indirect mechanisms impacting the cellular glutamate pool. Genetic analysis previously linked MT123 to a population of *Castellaniella* that bloomed—concurrent to nitrate removal—during a biostimulation effort to reduce groundwater nitrate concentrations at MT123’s location of origin. Thus, our analyses provide novel insight into mechanisms of acclimation to acidic conditions in a strain with significant potential for nitrate bioremediation.

**IMPORTANCE:** Nitrate pollution in groundwater is a serious threat to both environmental and human health. This nitrate pollution can come from a variety of sources including fertilizers, sewage, and industrial discharge. Certain bacteria, known as “denitrifiers”, can convert this nitrate into harmless nitrogen gas, a process known as “denitrification”. Denitrifiers can be used to clean up nitrate-contaminated groundwater. However, their ability to do this can be disrupted by changing environmental conditions. For example, groundwater that is polluted with nitrate is often acidic. Acidic conditions make it challenging for denitrifiers to survive, which results in less conversion of nitrate to nitrogen gas. In this study, we investigated how one denitrifying bacterium—originating from acidic, nitrate-contaminated groundwater—can cope with acidic conditions

## INTRODUCTION

Nitrate pollution in groundwater is a major environmental health hazard, globally (1). This nitrate originates from sewage runoff, livestock facilities, septic systems, industrial wastewater, agricultural fertilization, and atmospheric deposition (2). At concentrations as low at 50 mg/L, nitrate ingestion can cause methemoglobinemia in neonates and young children (2). Groundwater nitrate concentrations exceeding this threshold have been reported across the world (3).

Denitrifying bacteria have been leveraged for nitrate removal in groundwater (4). Biostimulation of contaminated groundwater with organic carbon promotes rapid proliferation of denitrifying bacteria, reducing nitrate to dinitrogen gas (5–8). However, these efforts are challenged by additional dynamic physiochemical conditions to which these bacteria are responsive. For example, co-occuring contaminants like heavy metals can inhibit the activities of enzymes in the denitrification pathway (9). Also, acidification is a common co-occurring problem in nitrate-contaminated groundwater (10), where the presence of the nitrate can itself promote acidification (13–15). Mildly acidic conditions with pHs lower than 6.0-6.5 inhibit the activity of nitrous oxide reductase NosZ (16–18) and decrease activity of nitrite reductases (19). Inhibition of individual steps within the denitrification pathway leads to the undesirable consequence of partial denitrification—resulting in the release of reactive nitrogen oxides such as nitrite, nitric oxide, and nitrous oxide (20,21). Thus, it is critical to understand how denitrifiers respond to these common co-occurring environmental stressors.

The Oak Ridge Reservation (ORR, Oak Ridge, TN, USA) is a well-characterized experimental site for examining the ecological impacts of legacy industrial waste (22). Previously, industrial waste generated from nuclear processing operations was dumped into four unlined waste ponds at ORR, known as the S-3 ponds. The soils and groundwater near the point source are highly acidic due to high concentrations of uranium nitrate in the industrial waste (22). The genus *Castellaniella* are facultatively anaerobic, denitrifying bacteria (23) that are highly abundant in the contaminated soils and groundwater at ORR (24). Native *Castellaniella* strains were found to be important for nitrate remediation in contaminated ORR soils (25), and *Castellaniella* was the dominant genus that bloomed following a successful biostimulation campaign that decreased local contaminating nitrate concentrations (25). In a previous analysis, we reported that this genus was commonly found in global anthropogenically impacted sites including contaminated soil and water, wastewater treatment plants, and the built environment (24).

We isolated *Castellaniella* sp. str. MT123 (MT123) from highly contaminated ORR groundwater taken from within the S-3 ponds contamination plume (26). This strain is a representative of a highly abundant and persistent operational taxonomic unit (OTU) that was first reported at the site following biostimulation efforts in the early 2000s (24). In a sediment core collected after biostimulation with ethanol and bicarbonate, this OTU was reported at 70% relative abundance (25). MT123 grows optimally under the mildly acidic conditions of pH 5.5. The strain is a complete denitrifier, retaining this activity even at lower pH (24). We hypothesized that this strain must have robust mechanisms for maintaining intracellular pH during growth under mildly acidic conditions. Here, we characterized the mechanisms of pH homeostasis in MT123. RB-TnSeq analysis was used to identify single gene mutants with decreased fitness at pH 5.5 compared to 7.5. We paired these results with proteomic and metabolomic analyses to characterize systems-level changes occurring during mildly acidic growth conditions. Our results show contrasting roles of glutamate and branched chain amino acids (BCAA) in the cellular response to mildly acidic growth conditions.

## MATERIALS AND METHODS

### Media and Culture Conditions

As described previously, *Castellaniella* sp. str. MT123 was isolated from groundwater samples taken from the contaminated well FW104 near the former S-3 waste ponds at the ORR (26). Glycerol stocks (10%) of MT123 were streaked on R2A agar plates (per liter: 0.5 g casein hydrolysate, 0.5 g dextrose, 0.5 g soluble starch, 0.5 g yeast extract, 0.3 g dipotassium phosphate, 0.3 g sodium pyruvate, 0.25 g proteose peptone, 0.25 g meat peptone, 15 g agar, and 0.05 g MgSO_4_7H_2_O). For pre-cultures, 5 mL of R2A broth was inoculated with five individual colonies from the R2A agar plates and grown at 30°C aerobically with shaking (200 rpm). For experimental work, the overnight preculture was inoculated in Modified UGA Medium. The Modified UGA Medium contained, per liter, 0.6 g NaH_2_PO_4_, 20 mL of 1 M sodium lactate, 40 mL of a 25X UGA Salts stock solution, 1 mL of a 1000X DL Vitamins stock, and 10 mL of a 100X DL minerals stock. The 25X UGA Salts stock solution contained, per liter, 250 mg NaCl, 367 mg CaCl_2_ · 2H_2_O, 12.32 g MgSO_4_ · 7H_2_O, 2.5 g KCl, and 5 g NH_4_Cl. The 1000X DL Vitamins stock and the 100X DL minerals stock recipes are described previously (27). For any cultures grown in anaerobic conditions, the medium also contained 10 ml of 1 M NaNO_3_ per liter with an 80%/20% N_2_/CO_2_ headspace. The pH of the medium was adjusted as indicated - 4.5 to 7.5 - with HCl or NaOH. Different buffers were used based on the desired culture pH. For cultures which were grown in pH<7, we added 7.8 g of MES buffer per liter. For cultures which were grown in pH≥7, we added 8.4 g of MOPs per liter.

### Growth Phenotyping Experiments

Growth experiments were performed in 100-well plates in a Bioscreen incubating plate reader (Growth Curves Ltd.) with a final volume of 400 µL of the Modified UGA Medium. For growth under denitrifying conditions, the Bioscreen was placed in an anaerobic chamber under a 78% N_2_ / 20% CO_2_ / 2% H_2_ headspace. For the amino acid additions, all amino acids were added at 1 mM concentrations. The Bioscreen monitored growth by optical density measurements at 600 nm (OD600) once every hour. The Bioscreen was set to shake cultures continuously at low amplitude throughout each growth experiment while holding the temperature at 30 °C.

### Genomic Analysis

The MT123 genome was published previously (24) and is available under BioProject number PRJNA1100609 on NCBI. The protein annotations used for this study are described in **Table S1**. Between the time of analysis and writing of this report, the completed version of the MT123 genome described above became publicly available. **Table S2** relates the locus tags used for this study to the current version of the MT123 genome. Proteins previously identified as being involved in acid tolerance in *Escherichia coli* K-12 or *Helicobacter pylori* strain J99 were utilized to perform a BLASTp search against the MT123 genome to identify homologs (28). Protein sequences were retrieved from UniProt.

### RB-TnSeq Experiments

The RB-TnSeq mutant library for MT123 was constructed by insertion of a barcoded mariner transposon as described previously (29). Expanded methodology can be found in the **SUPPORTING INFORMATION.** The individual mutants are mapped to the MT123 genome based on the annotations provided in **Table S1**. **Table S2** relates the locus tags used for this study to the current version of the MT123 genome.

A glycerol stock (∼1 mL) of the MT123 mutant library was inoculated into and grown aerobically in 100 mL R2A medium with 10 μg/mL kanamycin. These precultures were grown at 30°C and shaken at 200 rpm. Precultures were grown to approximately mid-log phase (OD600 = 0.5) and 1 mL of 1.0 OD600 equivalent of preculture was recovered for sequencing to use as the reference library for all challenge condition cultures. Samples were spun in a microcentrifuge at 12,000 x g for 3 minutes. The supernatant was removed, and cell pellets were stored at -80°C for sequencing. Additionally, 1 mL of 0.5 OD600 equivalent of preculture was inoculated into 10 mL of experimental medium.

Experimental cultures were grown in five replicates in the modified UGA medium described above using four different challenge conditions: aerobic (*i.e.,* normal atmospheric conditions) at pH 5.5, aerobic at pH 7.5, anaerobic at pH 5.5, and anaerobic at pH 7.5. Experimental cultures were grown at 30°C. The anaerobic cultures were grown in sealed hungate tubes with an 80%/20% N_2_/CO_2_ headspace. Aerobic cultures were grown in capped test tubes, shaken at 200 rpm. Experimental cultures were recovered at approximately mid-log phase (OD_600_ ≈ 0.5), where 1 mL of 1.0 OD600 equivalent was harvested and spun down in a microcentrifuge at 12,000 x g for 3 minutes. The supernatant was removed, and cell pellets were stored at -80°C prior to genomic DNA extraction, PCR amplification of the DNA barcodes, and barcode sequencing (BarSeq) (29).

### RB-TnSeq Data Processing

We calculated gene fitness scores using a previously described approach (29). Averages of strain fitness values for each challenge condition were first calculated to give a gene fitness value, after which several *Δfitness* values of interest were calculated. *Δfitness* values were calculated by subtracting the gene fitness value of the base condition (pH 7.5) from the challenge condition (pH 5.5). Genes with a large positive (*Δfitness* >1) or large negative (*Δfitness* <-1) change in fitness value were selected for further analysis (30).

### Growth Conditions for Proteomic and Metabolomic Analyses

Five individual colonies of wild-type MT123 were inoculated from an R2A agar plate into 10 mL of R2A growth medium, which was then grown overnight as described above. A 40-fold dilution of the pre-culture was performed into 50 mL of modified UGA medium, described above. Cultures for proteomic analyses were performed in triplicate, and cultures for metabolomic analyses were performed in sextuplicate. These experiments utilized the same four challenge growth conditions as the RB-TnSeq analysis described above. All experimental growth media were inoculated with 500 μL of preculture. The anaerobic experimental cultures were grown in sealed 100 mL serum bottles with an 80%/20% N_2_/CO_2_ headspace. Aerobic experimental cultures were grown in 250 mL Erlenmeyer flasks and shaken at 200 rpm. All cultures were grown at 30°C. When the cultures reached an OD_600_ of 0.5, each culture was spun in a centrifuge at 4,000 rpm for 5 minutes at 25°C. The supernatant was removed, and the pellet was washed with 5 mL of a pH 7.4 PBS buffer (per liter: 0.8 g NaCl, 0.02 g KCl, 0.144 g Na_2_HPO_4_, 0.024 g KH_2_PO_4_). Washed cell pellets were stored at -80°C prior to metabolomic and proteomic analyses.

### Proteomic Analysis

Expanded methodology for the proteomic analyses can be found in the **SUPPORTING INFORMATION** file. Briefly, protein was extracted from cell pellets and tryptic peptides were prepared by following established proteomic sample preparation protocol (31). The resulting peptide samples were analyzed on an Agilent 1290 UHPLC system coupled to a Thermo Scientific Orbitrap Exploris 480 mass spectrometer for discovery proteomics (32). Briefly, peptide samples were loaded onto an Ascentis® ES-C18 Column (Sigma–Aldrich, USA) for chromatographic separations. Eluting peptides were introduced to the mass spectrometer operating in positive-ion mode and were measured in data-independent acquisition (DIA) mode. DIA raw data files were analyzed by an integrated software suite DIA-NN (33). The databases used in the DIA-NN search (library-free mode) is the protein FASTA sequences based on the MT123 genome annotations provided in **Table S1,** plus the protein sequences of common proteomic contaminants. **Table S2** relates the locus tags used for this study to the current version of the MT123 genome. Output main DIA-NN reports were filtered with a global false discovery rate set at 0.01 (FDR <= 0.01) on both the precursor level and protein group level. The Top3 method, which is the average MS signal response of the three most intense tryptic peptides of each identified protein, was used to quantify proteins in the samples (34,35). Clusters of Orthologous Genes (COGs) categories were assigned to the MT123 proteome using reCOGnizer (36).

### Metabolomic Analysis

Frozen cell pellets were resuspended in 500uL water, frozen at -80C, lyophilized and then homogenized with 3.2mm stainless steel beads (BioSpec Products) in a Mini Bead Beater (BioSpec Products) three times for 5 seconds each with 10 seconds cool down in between; homogenized cell material was stored at -80C. On the day of LC-MS/MS analysis, samples were resuspended in 150uL of methanol containing internal standard mix **(Table S4**). The extracts were vortexed twice for 10 seconds, bath sonicated in ice water for 15 minutes, centrifuged (10,000 rcf, 5 min, 10°C) and then supernatant was filtered via 0.22um PVDF microcentrifuge filtration (10,000 rcf, 5 min, 10°C). Filtrate was transferred to amber glass LC-MS/MS vials for analysis. Briefly, metabolites were analyzed by both reverse phase chromatography and hydrophilic interaction chromatography each followed by tandem mass spectrometry analysis. Metabolite separations were performed using an Agilent 1290 LC stack and detected using an Thermo Q Extractive hybrid quadrupole-Orbitrap mass spectrometer. LC-MS/MS parameters are provided in **Table S4**. Internal and external standards as well as extraction and injection controls were used for quality control purposes.

### Metabolomic Data Processing

Raw datafiles are available in MZML format through https://gnps2.org/status?task=29f23392a1414aa2a75002938e2fecec. Metabolites were annotated using Metatlas (https://github.com/biorack/metatlas) (37) to compare m/z, retention time and fragmentation spectra from samples to reference standards analyzed using the same methods. Annotations are provided in **Table S3.** To identify significant changes in metabolome abundance, a Welch’s t-test was performed to compare metabolite abundances at pH 5.5 to those at pH 7.5 under both growth conditions. All p-values were corrected for multiple comparisons using the Benjamini-Hochberg method.

## RESULTS

### MT123 grows optimally under mildly acidic conditions

Concentrations of dissolved oxygen in groundwater can fluctuate over seasonal and storm-event timescales (38,39). Thus, we first examined the growth of MT123 under both aerobic (i.e., normal atmospheric conditions) and denitrifying growth conditions at both neutral (pH 7.5) and mildly acidic (pH 5.5) pHs. Across the ORR subsurface, groundwater samples have been recorded in a wide range of pH, from 3 to 10.5 (40). pH 5.5 was chosen for this study as it reflects the pH of ORR groundwater well FW104, the isolation site of MT123. Consistent with our prior findings (24), under aerobic growth conditions, a higher maximal growth rate and slightly higher carrying capacity was observed at pH 5.5 relative to 7.5 **(Fig. 1A)**. Under denitrifying conditions, a greater proportion of the resulting nitrite is in the form of free nitrous acid (FNA), a potent protonophore (41,42) that is inhibitory to denitrifiers (43). Nonetheless, MT123 retains robust growth at pH 5.5 **(Fig. 1B)**.

**Figure 1.**
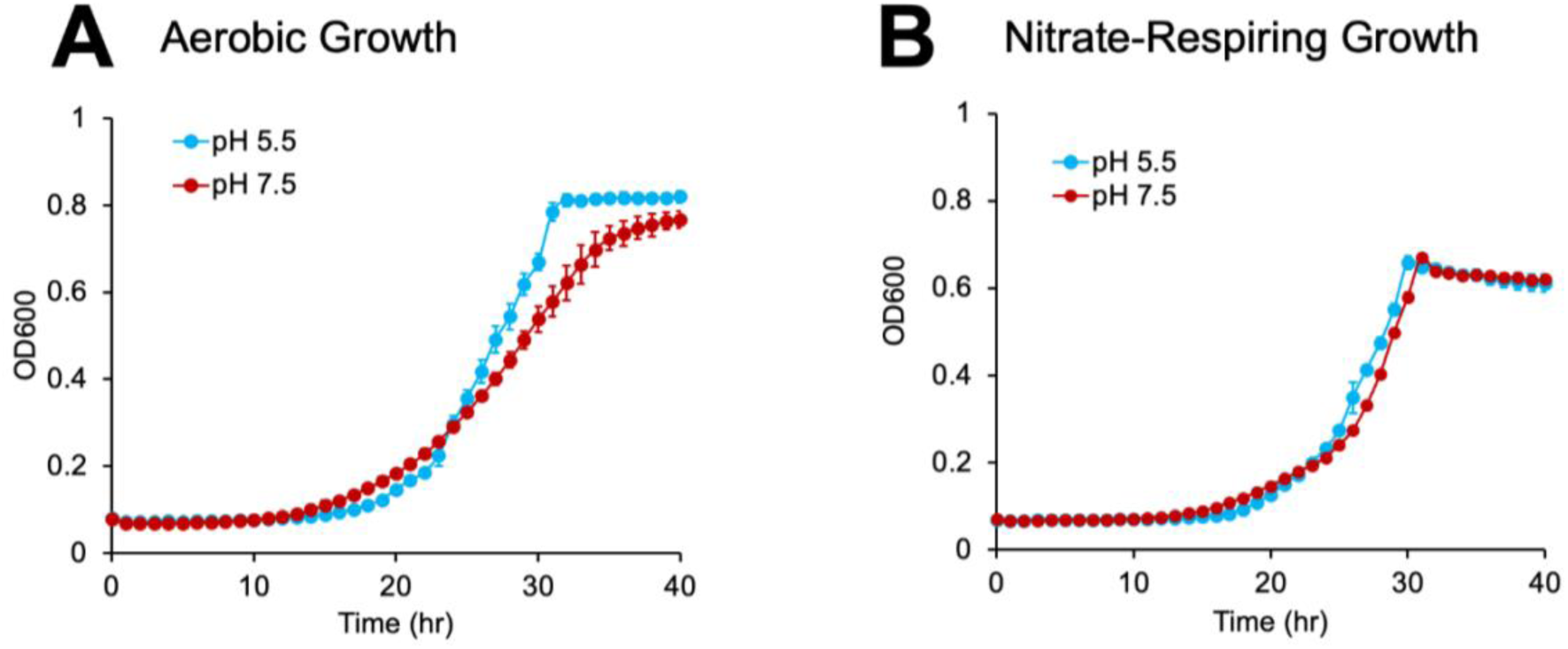
Growth of MT123 at pH 5.5 and 7.5 (legend shown on figure) under aerobic (A) and denitrifying (B) conditions. Data points represent the average of triplicate experiments. Error bars represent ±SD.

### Genomic analysis of MT123 reveals a lack of classic acid tolerance systems

In many bacteria, acid tolerance under mildly or moderately acidic conditions depends on the cytoplasmic buffering activity of amino acid decarboxylase enzymes (namely glutamic acid, arginine, lysine, and ornithine decarboxylases) paired with an amino acid antiporter systems to exchange extracellular amino acids with the decarboxylation products (44,45) **(Fig. 2).** Analysis of the MT123 genome revealed the strain lacked the glutamic-acid dependent system **(Fig. 2A)** and had only partial versions of the other three systems **(Fig. 2B-C).** The MT123 genome encodes a putative arginine/lysine/ornithine decarboxylase, which may carry out the decarboxylation of some or all three amino acids. However, the genome lacks the genes encoding the associated antiporters AdiC (arginine/agmatine antiporter), CadB (cadaverine/lysine antiporter), and PotE (putrescine/ornithine antiporter). Another acid tolerance system involves arginine deamination to citrulline. This is followed by conversion of citrulline to ornithine and carbamoyl phosphate. Ornithine is exchanged for arginine by an antiporter and the carbamoyl phosphate is decarboxylated by carbamate kinase to yield ammonium. MT123 encodes the enzyme (ArgF, ornithine carbamoyltransferase) for the second step in the pathway; however, it lacks a homolog for the arginine deaminase ArcA as well as the ArcD antiporter and the carbamate kinase AllK.

**Figure 2.**
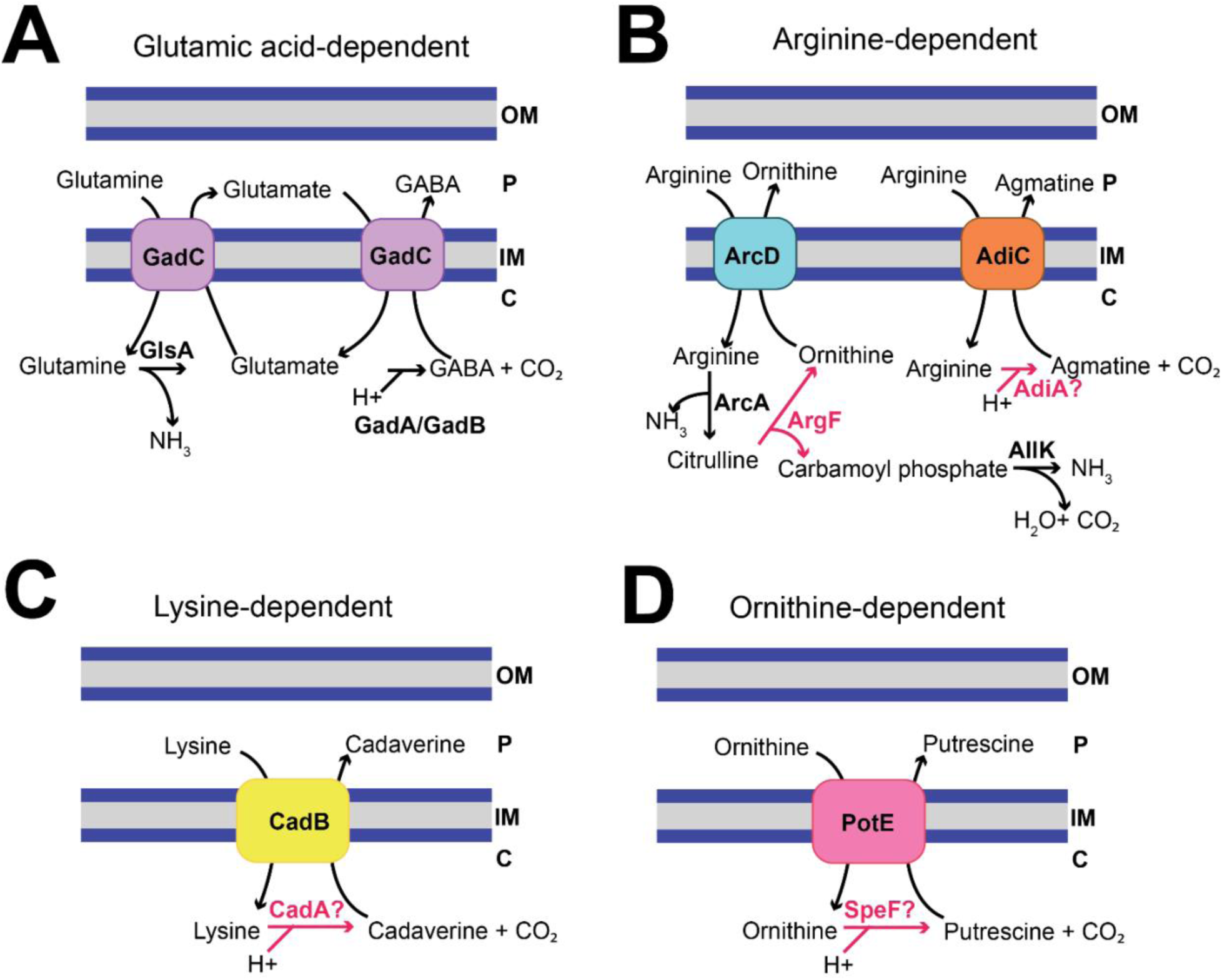
Analysis of the MT123 genome for known acid tolerance mechanisms under mildly and moderately acidic conditions. All four shown systems utilize an amino acid decarboxylase coupled to an antiporter for exchange of the decarboxylase substrate and product. Only the enzymatic steps highlighted in red are predicted in MT123 from the genome analysis. **(A)** Glutamic acid-dependent pathway. GlsA = glutaminase, GABA = gamma-aminobutyric acid, GadC = glutamate/GABA antiporter, GadA = glutamate decarboxylase A, GadB = glutamate decarboxylase B. **(B)** Arginine-dependent pathway. ArcD = arginine/ornithine antiporter, ArcA = arginine deiminase, ArgF = ornithine carbamoyltransferase, AllK = carbamate kinase, AdiC = arginine/agmatine antiporter, AdiA = arginine decarboxylase. **(C)** Lysine-dependent pathway. CadA = lysine decarboxylase, CadB = cadaverine/lysine antiporter. **(D)** Ornithine-dependent pathway. SpeF = ornithine decarboxylase, PotE = putrescine/ornithine antiporter. Note that the genome has a decarboxylase annotated as an arginine/lysine/ornithine decarboxylase that is a homolog to AdiA, CadA, and SpeF.

At very low pH, acid-tolerant bacteria also utilize the urease system (encoded by the *ureIABEFG* operon) to produce alkali in the form of ammonia (46). However, this system is absent in the MT123 genome. Proteins previously associated with acid tolerance responses (44) that are encoded in the MT123 genome include RecA (repair and maintenance of DNA damage), DnaK (chaperone Hsp70), DnaJ (chaperone Hsp40), ClpXP (protease), ClpAP (protease), GroEL (chaperone Hsp60), GroES (chaperone Hsp10), and UvrABC (excinuclease). We next examined if any of these identified systems were involved in the MT123 growth at mildly acidic pH or if the cells utilize a yet-undescribed system for acid tolerance.

#### Alteration of amino acid metabolism during growth at pH 5.5

We compared the proteomic response of cells grown at pH 5.5 to cells grown at pH 7.5 under both aerobic and (anaerobic) denitrifying conditions. Differentially abundant proteins at pH 5.5 were identified through comparison to the proteome of cultures grown at pH 7.5 (notated as Log2FC). When considering both the aerobic and denitrifying growth conditions, we detected 485 significantly (*adjusted p<0.05*) differentially abundant proteins at pH 5.5 **(Tables S5-S8).** However, the proteomic response to growth at pH 5.5 was more pronounced under aerobic conditions (432 differentially abundant proteins) compared to denitrifying conditions (158 differentially abundant proteins). Notably, none of the putative acid tolerance proteins identified in our genome analysis changed significantly in abundance at pH 5.5 under either growth condition.

We assigned COG categories to the entire predicted MT123 proteome as well as both the aerobic and anaerobic differentially abundant proteomes. Under aerobic conditions, the distribution of COG categories across the entire MT123 proteome and the differentially abundant proteome at pH 5.5 were significantly enriched in COG category E (22% vs. 15%, *Fisher’s exact test, adjusted p = 0.002*), that encompasses amino acid transport and metabolism **(Fig. 3).** Under denitrifying conditions, we observed a similar trend (24% vs. 15%) **(Fig. 3).** However, the statistical comparison following the FDR correction did not reach the threshold of significance due to the smaller number of differentially abundant proteins (*Fisher’s exact test, adjusted p = 0.06*). We observed that both glutamate metabolism and branched chain amino acid (BCAA) metabolism were significantly altered in the proteome at pH 5.5 under both growth conditions. Thus, we sought to examine the involvement of both sets of metabolic pathways in growth under mildly acidic conditions.

**Figure 3.**
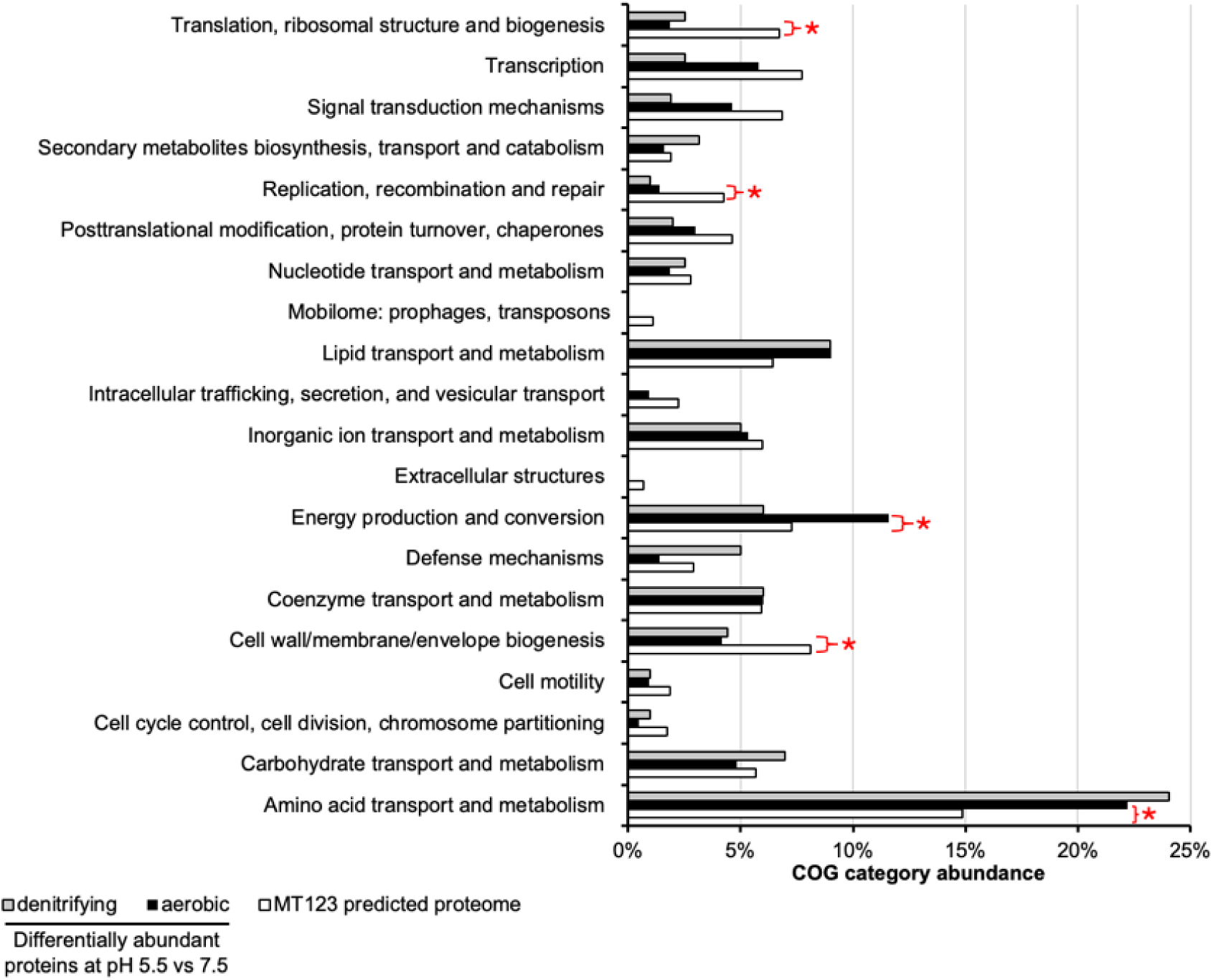
Abundances of the clusters of orthologous genes (COG) categories assignments for the differentially abundant proteins at pH 5.5 vs. 7.5 under aerobic (black bars) and denitrifying (grey bars) growth conditions. These data are shown alongside the distributions of COG category assignments for the entire MT123 proteome (white bars). A Fisher’s exact test was used to perform an enrichment analysis, comparing the entire proteome to the two differentially abundant proteomes. The Benjamini-Hochberg procedure was used to control for the false discovery rate (FDR). Red stars indicate an adjusted p-value <0.05.

#### Proteomic and metabolomic analysis suggests production of glutamate during pH 5.5 growth as an acid tolerance mechanism

As a general trend at pH 5.5, we observed an increase in abundance of enzymes that catalyze glutamate-yielding reactions and a decrease in abundance of enzymes that catalyze glutamate-consuming reactions **(Table 1).** For example, we saw increased abundance of the glutamate synthase large subunit (GltB) under both growth conditions (aerobic: +3.0 log2FC, denitrifying: +1.0 Log2FC) and the glutamate synthase small subunit (GltD) under aerobic growth (+3.3 log2FC), which are predicted to produce glutamate from glutamine and 2-oxoglutarate (alpha-ketoglutarate) **(Fig. 4A)**(47). Consistent with this observation, targeted metabolomic analyses revealed significantly decreased (-2.0 Log2FC) cellular glutamine content at pH 5.5 relative to pH 7.5 under aerobic conditions **(Fig 4B, Table S9).** In contrast, the glutamate dehydrogenase (GdhA)—which is predicted to consume glutamate, producing ammonium and 2-oxoglutarate(48)—decreases in abundance under aerobic growth conditions (-2.6 log2FC). Glutamate is also converted to 2-oxoglutarate through the action of 4-aminobutyrate aminotransferase PuuE in reaction with either succinate semialdehyde or glutarate semialdehyde. At pH 5.5, we observed decreased abundance of PuuE under aerobic growth conditions (-6.9 log2FC). These proteomic data suggest decreased cellular levels of 2-oxoglutarate at pH 5.5 relative to pH 7.5. However, we could not measure 2-oxoglutarate levels accurately due to challenges with isomer resolution.

**Figure 4.**
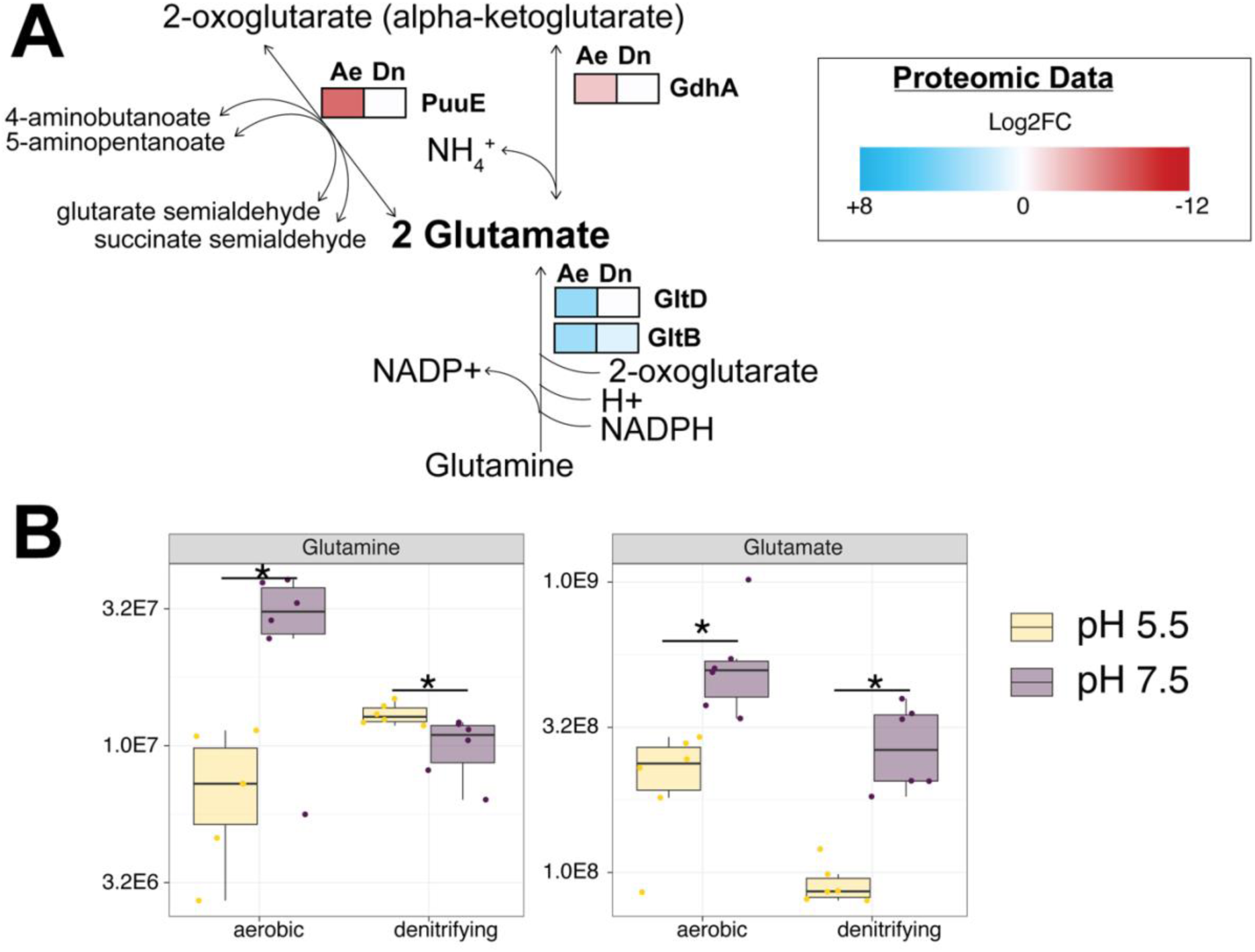
Integrated proteomic and metabolomic insights into the role of glutamate metabolism in the cellular response to growth under mildly acidic conditions. (A) Patterns of protein abundances involved in glutamate metabolism. Heat maps display the average (n=3) Log2FC values for the individual proteins under aerobic (Ae) and denitrifying (Dn) growth conditions. The heat map scale (displayed as Log2FC abundances at pH 5.5 vs. 7.5) is at the bottom of the panel. Blue boxes indicate increased protein abundance (p<0.05), red boxes indicate proteins that were significantly decreased in abundance (p>0.05), and white boxes indicate no significant difference in protein abundance. Statistical comparisons were performed with a Welch’s t-test using the Benjamini-Hochberg procedure to control for the false discovery rate (FDR). (B) Relative abundances of intracellular glutamine and glutamate. Individual data points represent replicate experiments (n = 6). Statistical comparisons were performed with a Welch’s t-test using the Benjamini-Hochberg procedure to control for the false discovery rate (FDR). Asterisks (*) indicate an adjusted p-values <0.05.

**Table 1.**
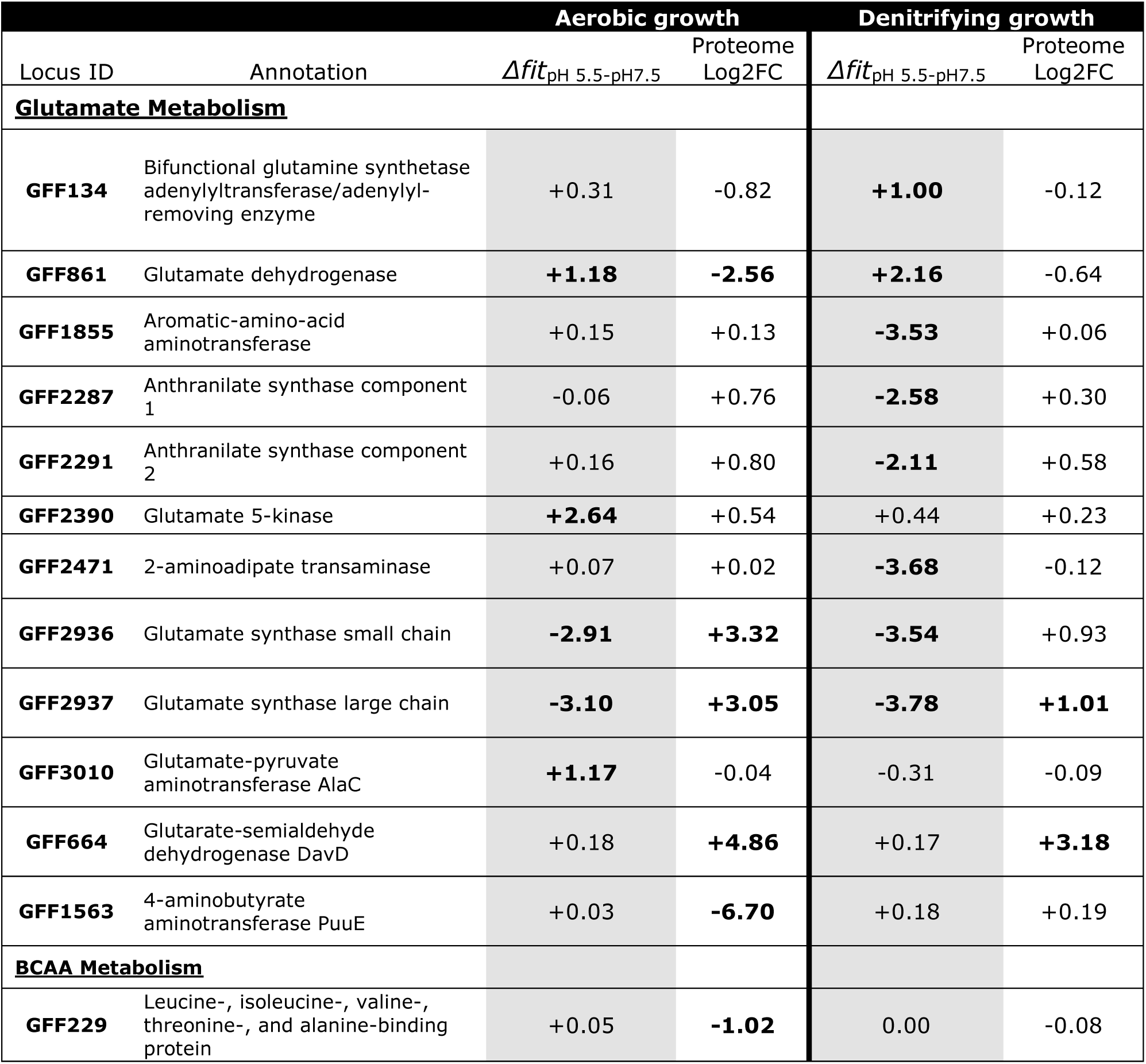
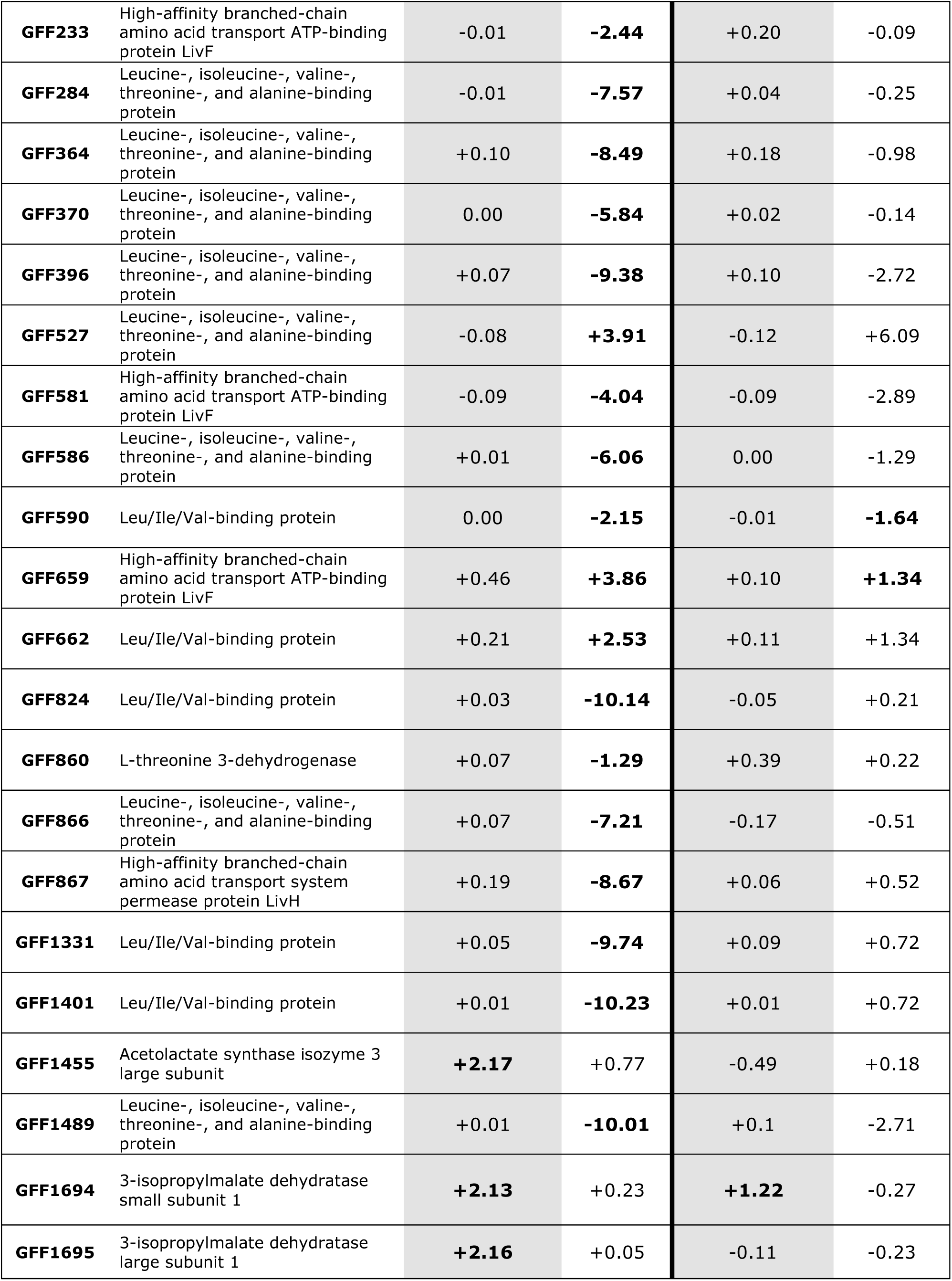
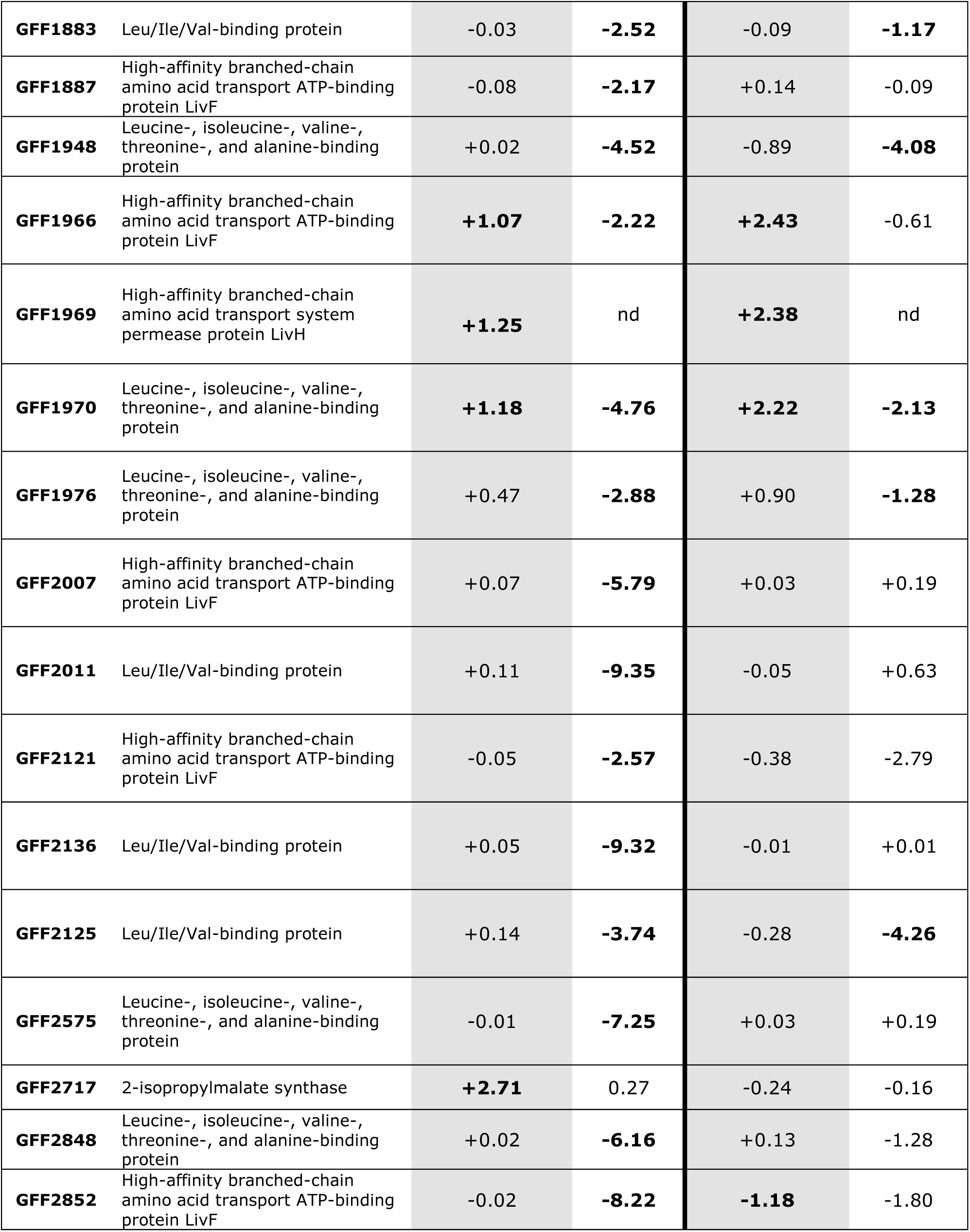
Genes involved in glutamate metabolism or BCAA metabolism with large fitness changes in pH 5.5 compared to pH 7.5. Δfit_pH5.5-pH7.5_ values represent fitness change values between growth at pH 5.5 and pH 7.5, calculated from RB-TnSeq data. Significant fitness changes are bolded and defined as |log_2_| ≥ 1. Accompanying Log2FC values represent fold changes from proteomics data, with significant fold change being in bold and defined as |log2FC| ≥ 1 and a p-value < 0.05. The linked RefSeq locus tags can be found in **Table S2**.

At pH 5.5, intercellular glutamate is decreased relative to pH 7.5 under both growth conditions **(Fig. 4B, Table S9).** This is despite the increased abundance of proteins involved in glutamate production and decreased abundance of proteins involved in glutamate consumption. Thus, growth at pH 5.5 appears to increase the cellular demand for glutamate, likely triggered by this glutamate depletion. While both the metabolomic and proteomic data were collected at the same points in MT123 growth, the two data types inherently represent “snapshots in time” of different scales.

The turnover of metabolites can occur at orders of magnitude faster rates than the turnover of proteins (49–51). The fact that we simultaneously observe increased abundances of proteins involved in glutamate production alongside decreased cellular glutamate abundance indicates consistent and on-going high rates of glutamate consumption within the cell during growth at pH 5.5.

Among the proteins that decreased in abundance during growth at pH 5.5, perhaps the most striking trend was the large number of BCAA transporters. When analyzing the MT123 genome, we observed a remarkably high number of BCAA transporters including 25 LivH/LivM homologs (the permease subunit). In *E. coli* K12 the Liv transporter is a common transporter for L-leucine, L-isoleucine, L-valine, and the BCAA precursor L-threonine, although their specificity may vary among species (52–55). A multiplicity of BCAA transport systems has been reported in other bacteria, likely due to the critical functions of BCAAs as key metabolic intermediates and in signaling of cellular metabolic status (53,55,56). However, we failed to find a report of a bacterial genome with as many BCAA transporter homologs as MT123. In the pH 5.5 proteomes, we observed decreases in abundance of seven homologs of the high-affinity BCAA transporter ATP-binding subunit LivF; one homolog of the high-affinity BCAA transporter permease subunit LivH; and 21 homologs of the BCAA binding protein of the Liv transporter **(Table 1)**, with a greater number of the transporters decreasing in abundance under aerobic growth conditions (29 proteins) compared to denitrifying conditions (6 proteins).

We next asked whether BCAAs accumulate in the cytoplasm during growth at pH 5.5—a possible signal to decrease the expression of BCAA importers. We observed significant increases in the abundance of valine at pH 5.5 under both aerobic and denitrifying conditions **(Fig. 5A, Table S9)**. We also observed significant increases in leucine and isoleucine abundances at pH 5.5 under aerobic growth conditions **(Fig. 5B, C, Table 8)**. While leucine and isoleucine abundances did increase at pH 5.5 under denitrifying conditions, this increase did not achieve the threshold of significance (**Fig 5A-C, Table S9**). No significant changes in abundances of the BCAA-precursor threonine were observed under either condition **(Fig. 5D, Table S9)**. Thus, we suggest that under mildly acidic conditions, intracellular concentrations of the BCAAs leucine, valine, and isoleucine increase, leading to repression of BCAA importers. In other gram-negative bacteria, leucine, valine, threonine, and isoleucine are effectors of the global regulator Lrp (***L***eucine-responsive ***r***egulatory ***p***rotein)(57,58), with leucine-bound Lrp acting as a repressor of the *liv* operon in *E. coli* K-12 (59). MT123 encodes three Lrp homologs in its genome. We propose that the accumulation of BCAAs during growth under mildly acidic conditions leads to Lrp-mediated repression of its many *liv* operons.

**Figure 5.**
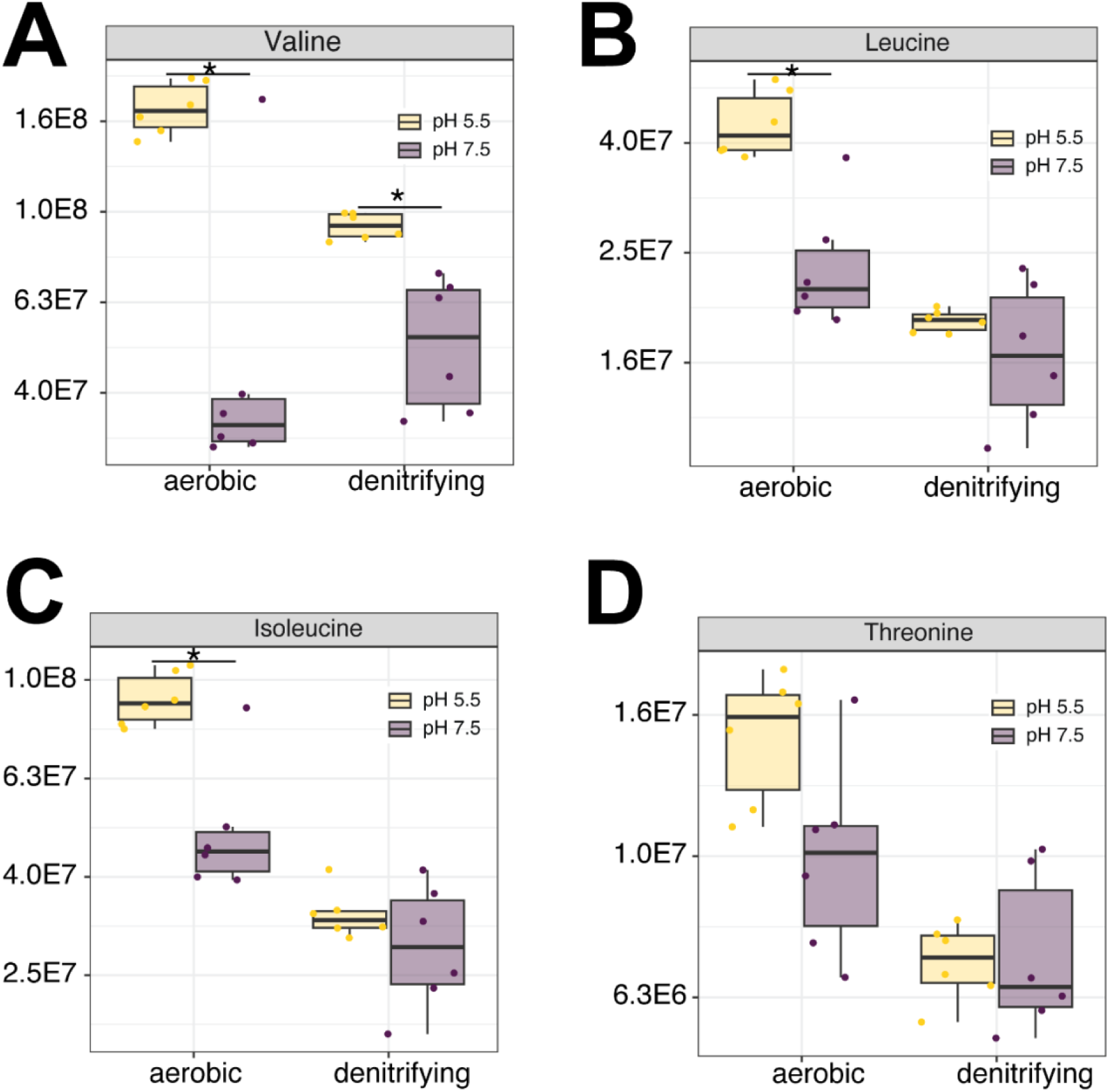
Changes in intracellular abundances of BCAAs and threonine during growth under mildly acidic conditions. Intracellular abundances of valine (A), leucine (B), isoleucine (C), and threonine (D). For all panels, individual data points represent replicate experiments (n = 6). Statistical comparisons were performed with a Welch’s t-test using the Benjamini-Hochberg procedure to control for the false discovery rate (FDR). Asterisks (*) indicate an adjusted p-values <0.05.

#### Opposing effects of glutamate and BCAAs on MT123 fitness during mild acid stress

While the proteomic and metabolomic data reveal the cellular response to mildly acidic conditions, these data do not imply whether these responses benefit the cells under these conditions. Thus, we constructed an RB-TnSeq fitness library to calculate fitness scores for 2,349 genes during growth at pH 5.5 (approximately 75% of MT123’s predicted protein coding genes). Changes in gene fitness values (*Δfitness*) were calculated from the difference between gene fitness values of pH 5.5 and pH 7.5 cultures **(Table S10).** Genes with large positive (*Δfitness* > 1) or large negative (*Δfitness* < -1) fitness changes (30) were analyzed further for involvement in either glutamate or BCAA metabolism. Negative fitness changes reflect gene disruptions that have a more detrimental effect on growth at pH 5.5 relative to pH 7.5. Positive fitness changes reflect gene disruptions that have a more beneficial effect on growth at pH 5.5 relative to pH 7.5.

The RB-TnSeq data support a model where glutamate accumulation is critical in the cellular response to growth at pH 5.5 **(Table 1).** Highly negative *Δfitness* values were observed for the glutamate synthase genes (*gltDB*) under both aerobic and denitrifying growth conditions (*Δfitness_aerobic_ =* -2.91 and -3.10; *Δfitness_denitrifying_ =* -3.54 and -3.78), consistent with our observation that these proteins increase in abundance at pH 5.5 **(Fig. 4, Fig. 6)**. The screen also uncovered additional genes predicted to be involved in glutamate production with negative *Δfitness* values that encode the 2-aminoadipate transaminase (*lysN*)(60), the aromatic amino acid aminotransferase (*tyrB*)(61), and the anthranilate synthase subunits 1 (*trpE*) and 2 (*trpD*)(62). As amino acid transaminases are readily reversible (63,64), we suggest that under the growth conditions utilized here that they might operate in the direction of glutamate synthesis **(Fig. 6).** In contrast, we observed that genes encoding enzymes predicted to be involved in glutamate consumption had positive *Δfitness* values, suggesting that glutamate consumption through these pathways is detrimental to cell fitness at pH 5.5 **(Table 1).** For example, the glutamate dehydrogenase *(gdhA)* gene had positive *Δfitness* values (*Δfitness_aerobic_ = +*1.18; *Δfitness_denitrifying_ = +*2.16). Consistent with this observation, the glutamate dehydrogenase (GdhA) decreased in abundance at pH 5.5 under aerobic growth conditions **(Fig. 6).** We also found that the glutamate 5-kinase gene (*proB*), which converts glutamate to glutamyl-5-phosphate(65), and the glutamine synthetase adenylyltransferase (glpE), which regulates the activity of the glutamine synthetase (GlnA; glutamate è glutamine)(66), had positive *Δfitness* under aerobic growth conditions **(Table 1).** In contrast to the other two transaminase-encoding genes, the *alaC* gene—encoding the glutamate-pyruvate aminotransferase(67)—had a positive Δ*fitness* under aerobic conditions, suggesting that it may be active primarily in the direction of 2-oxoglutarate and alanine formation from glutamate and pyruvate **(Fig. 6).**

**Figure 6.**
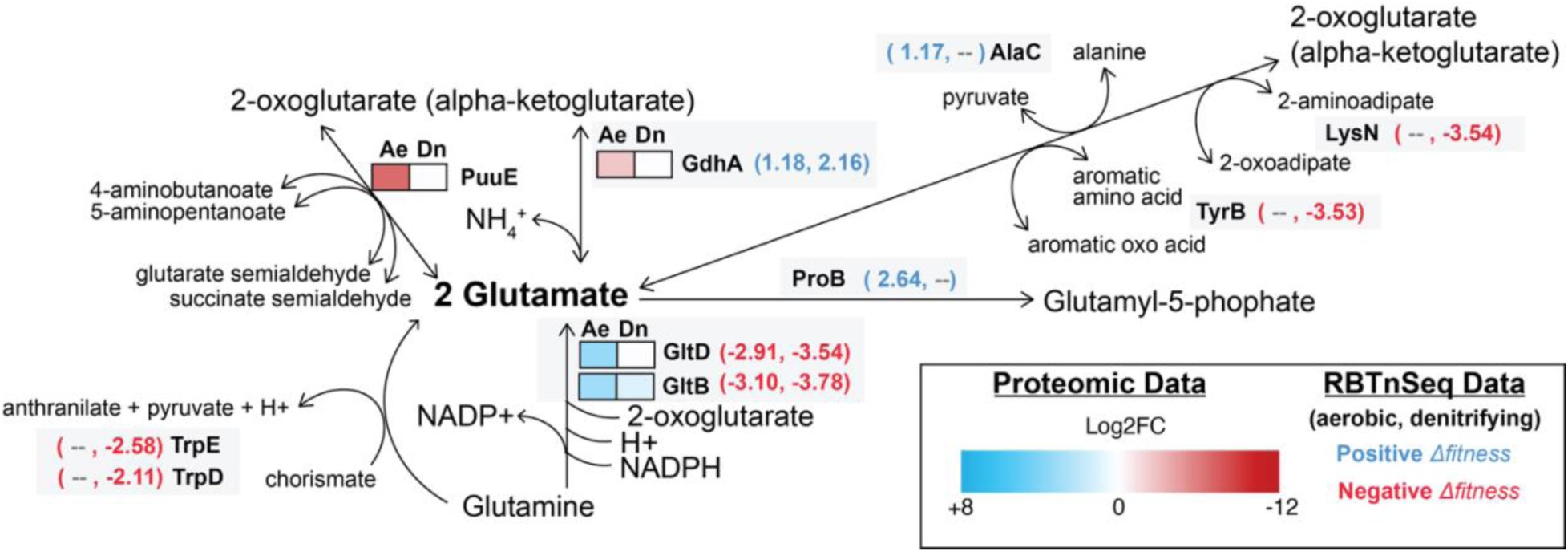
Integrated proteomic and mutant fitness assay (RB-TnSeq) insights into the role of glutamate metabolism in the cellular response to growth under mildly acidic conditions. Heat maps display the average (n=3) Log2FC values for the individual proteins under aerobic (Ae) and denitrifying (Dn) growth conditions. The heat map scale (displayed as Log2FC abundances at pH 5.5 vs. 7.5) is at the bottom of the panel. Blue boxes indicate increased protein abundance (p<0.05), red boxes indicate proteins that were significantly decreased in abundance (p>0.05), and white boxes indicate no significant difference in protein abundance. Statistical comparisons were performed with a Welch’s t-test using the Benjamini-Hochberg procedure to control for the false discovery rate. These data are overlaid with mutant fitness data for the gene encoding the enzyme. Genes with large positive (Δfitness > 1) or large negative (Δfitness < -1) are highlighted by the bracketed numbers next to the heat maps in either blue (positive) or red (negative) text. The first number is the Δfitness under aerobic conditions and second is under denitrifying conditions. If an enzyme lacks accompanying mutant fitness data, that means that there was no Δfitness under either growth condition.

These proteomic, metabolomic, and mutant fitness screen data suggest that intracellular glutamate is key for cellular growth at pH 5.5. We grew MT123 under both aerobic **(Fig. 7A)** and denitrifying **(Fig. 7B)** growth conditions at increasingly lower pHs until growth was eliminated (pH 4.5 for both conditions). We then grew the cells at pH 4.5, 5.0, 5.5, and 7.0 with an addition of 1 mM glutamate. Glutamate rescued any growth defects at pH 4.5 and 5.0 under aerobic conditions and at pH 5.0 under denitrifying conditions. Above pH 5.0, the growth improvement from the glutamate was minimal. While the MT123 genome does not encode a GadC homolog **(Fig. 2A)**, MT123 does seem to have the ability to import glutamate. We searched the MT123 genome for homologs to the *E. coli* GltS, GltP, and GltIJKL glutamate transporters. Putative homologs were identified for GltP (GFF2148) and GltIJKL (GFF926-929). Both GltP and GltIJKL have been linked to acid tolerance in other bacteria (68,69). Oddly, we observed decreases in the abundance of GltI (aerobic: -2.7 log2FC, denitrifying: -1.6 Log2FC) and GltJ (aerobic: -1.0 log2FC, denitrifying: -1.0 Log2FC) at pH 5.5 in our proteomics data. A second GltI homolog (GFF1522) was increased in abundance at pH 5.5 under aerobic conditions (+1.9 log2FC). However, the proteomics experiments were performed without glutamate supplementation where *de novo* synthesis is likely the major source of cytoplasmic glutamate pools rather than uptake—making it challenging to interpret these data in the context of our glutamate supplementation experiments.

**Figure 7.**
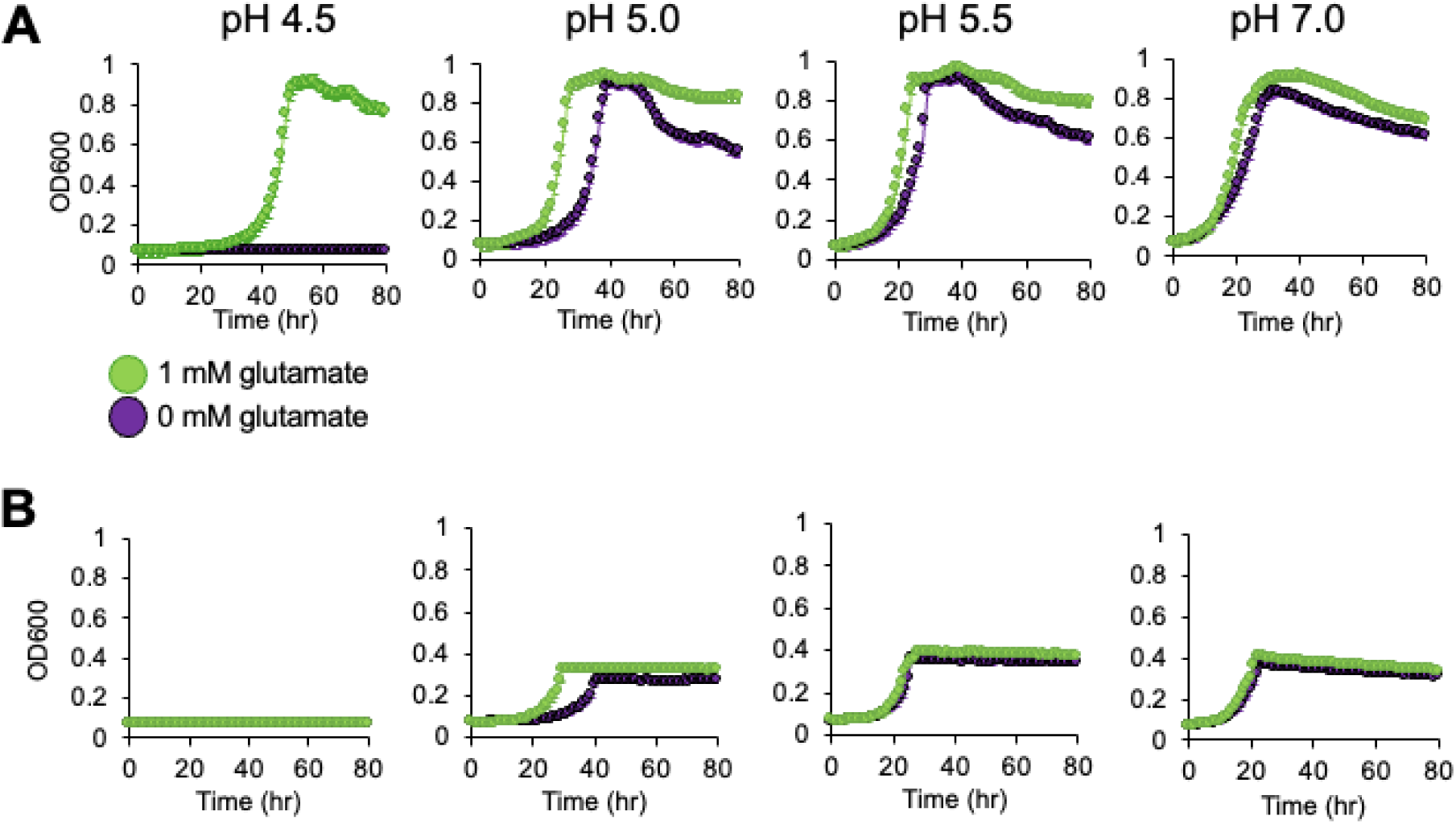
Growth of MT123 at varying pHs with 1 mM glutamate. Growth curves were performed under aerobic (A) and denitrifying (B) growth conditions. Individual points represent the average of three replicates. Error bars are present, though not always visible, and represent ±SD. The on-figure legend applies to all panels.

Given the negative *Δfitness* observed for *gltDB*, we considered that the conversion of glutamine to glutamate is the critical cellular activity conferring low pH tolerance, due to the consumption of a cytoplasmic proton in the process **(Fig. 6)**. Thus, we performed these same experiments with 1 mM glutamine. However, glutamine did not rescue growth at pH 4.5 and only slightly decreased lag time at pH 5.0 (**Fig. S1**). Taken together our results suggest that glutamate accumulation is critical for MT123 growth under mildly acidic conditions. We propose that the modulation of enzyme abundances **(Table 1)** to increase metabolite flux towards glutamate (and decrease its consumption via certain pathways) is an acclimatization response to mildly acidic conditions during both aerobic and denitrifying growth.

In contrast to glutamate, our mutant fitness data support a model where accumulation of one or more of the BCAA is detrimental to cellular growth at pH 5.5. While there is likely some degree of functional redundancy among the multiple BCAA transporters(53) in the MT123 genome, a cluster of genes encoding three of the Liv subunits (GFF1966, 1969, 1970) all had positive *Δfitness* values under both aerobic and anaerobic growth conditions **(Table 1).** This Liv transporter also decreased in abundance at pH 5.5 under both these growth conditions **(Table 1).** Consistent with our model, we also observed positive *Δfitness* values for several genes involved in BCAA biosynthesis. This included the gene encoding the acetolactate synthase large subunit (*ilvI, Δfitness_aerobic_ =* +2.17) which is part of both the isoleucine and valine biosynthesis pathways. Additionally, we observed *Δfitness* values for the 3-isopropylmalate dehydratase large (*leuC, Δfitness_aerobic_ =* +2.13; *Δfitness_denitrifying_ =* +1.22) and small subunits (*leuD, Δfitness_aerobic_ =* +2.16) as well as the 2-isotropylmalate synthase (*leuA, Δfitness_aerobic_ =* +2.71) —all part of the leucine biosynthesis pathway.

From these data, we hypothesized that accumulation of leucine, isoleucine, and/or valine—as was observed in our pH 5.5 metabolomics data **(Fig. 5)**—is inhibitory to cell growth at pH 5.5. We tested this hypothesis by repeating the same series of AA acid addition growth experiments performed above but with 1 mM additions of isoleucine leucine, valine, and threonine under both aerobic **(Fig. 8)** and denitrifying growth conditions (**Fig. 9**). Additions of isoleucine, leucine, or valine all inhibited growth at pH 5.0, with valine as the most inhibitory. These effects were more pronounced under denitrifying compared to aerobic growth conditions. In contrast, amendments with the BCAA precursor threonine had no impact on growth under the conditions tested. Considering these findings, we suggest that the Liv transporter (GFF1966, 1969, 1970) **(Table 1)** with positive *Δfitness* values is likely the primary transporter for leucine, isoleucine, and/or valine import by MT123 under the conditions studied here.

**Figure 8.**
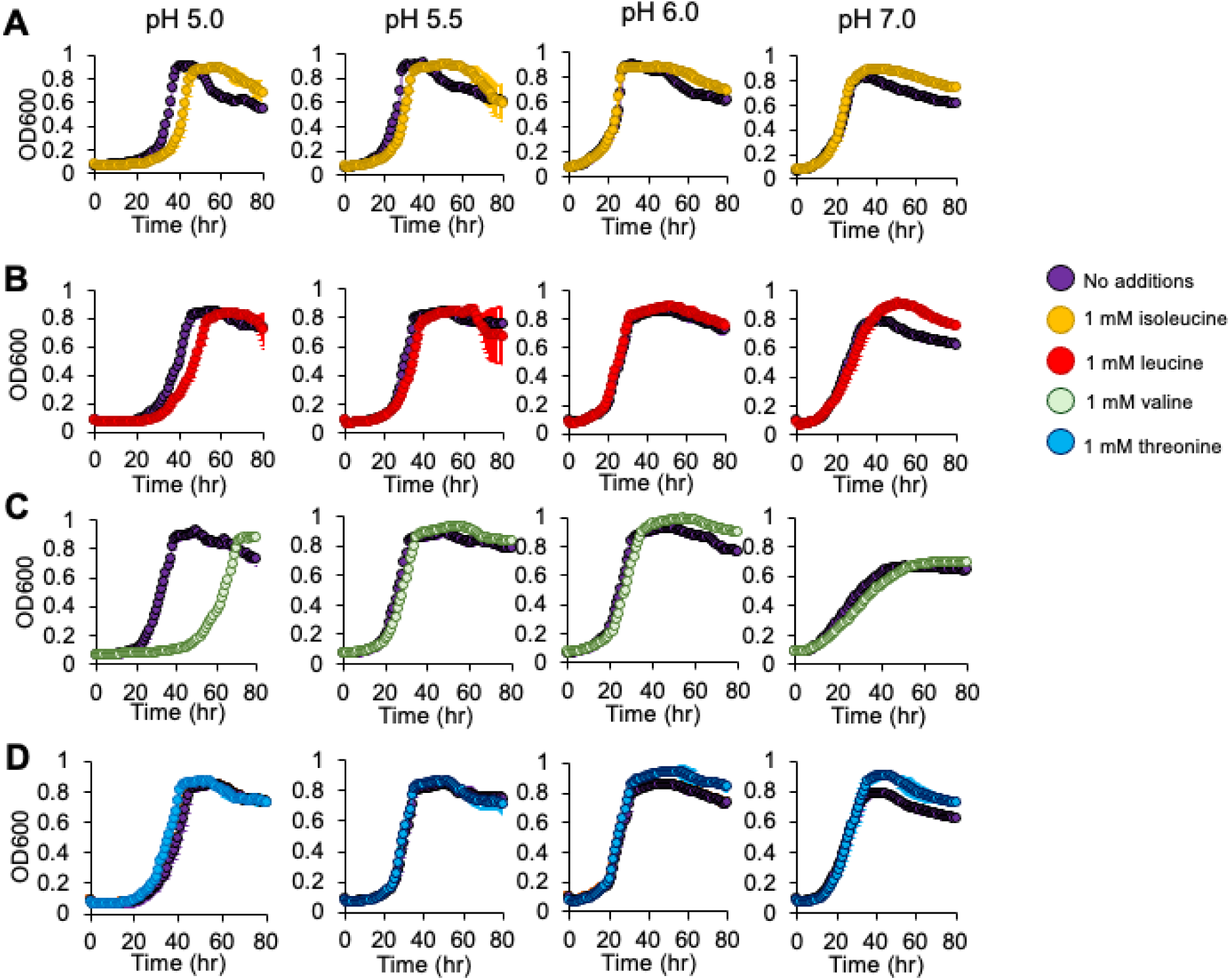
Growth of MT123 at varying pHs with 1 mM BCAA and threonine additions. Growth curves were performed under aerobic growth conditions. (A) Growth with 1 mM isoleucine. (B) Growth with 1 mM leucine. (C) Growth with 1 mM valine. (D) Growth with 1 mM threonine. Individual points represent the average of three replicates. Error bars are present, though not always visible, and represent ±SD. The on-figure legend applies to all panels.

**Figure 9.**
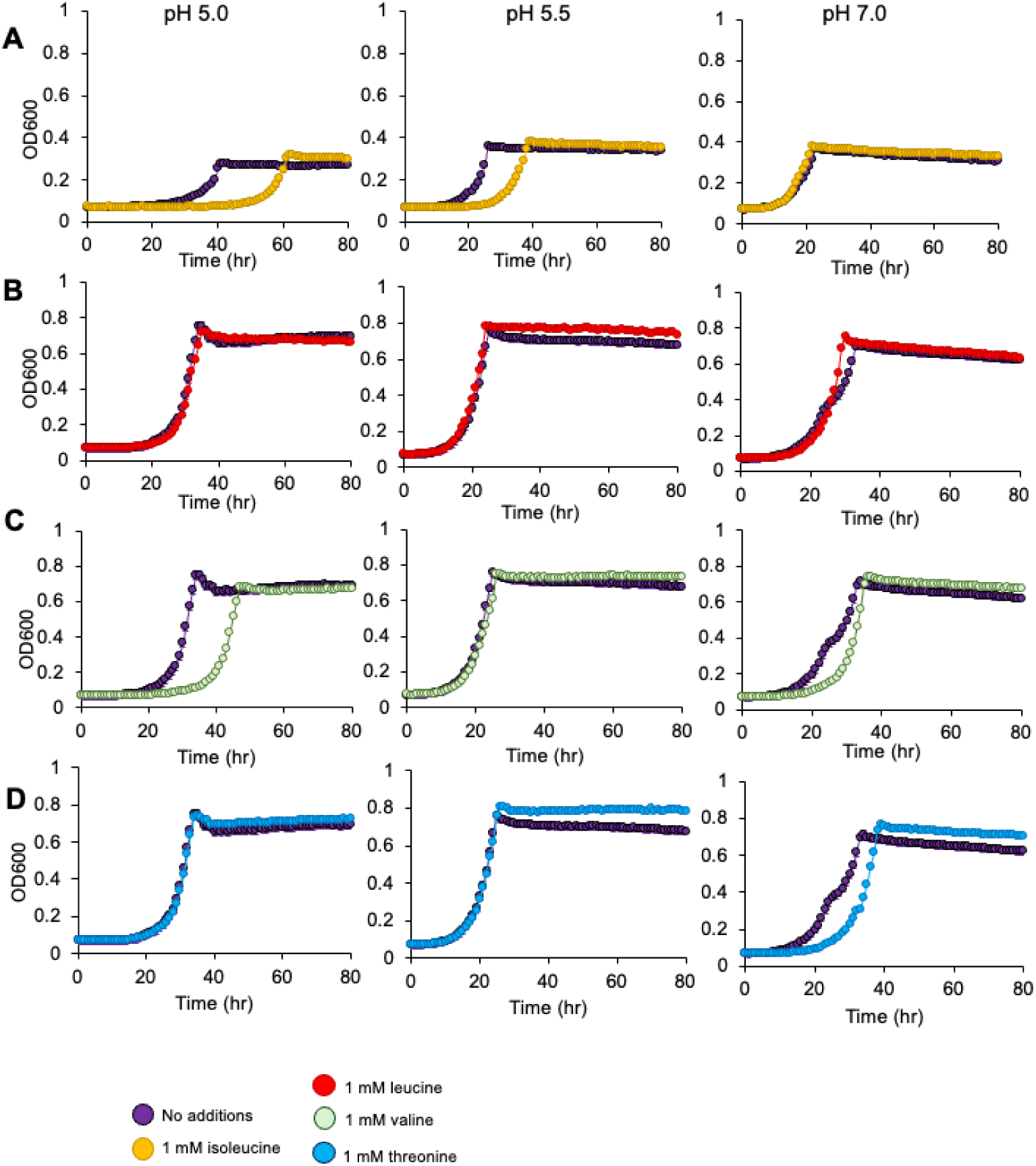
Growth of MT123 at varying pHs with 1 mM BCAA and threonine additions. Growth curves were performed under denitrifying growth conditions. (A) Growth with 1 mM isoleucine. (B) Growth with 1 mM leucine. (C) Growth with 1 mM valine. (D) Growth with 1 mM threonine. Individual points represent the average of three replicates. Error bars are present, though not always visible, and represent ±SD. The on-figure legend applies to all panels.

## DISCUSSION

While glutamate is often implicated in the bacterial response to mildly acidic growth conditions, MT123 lacks the canonical glutamate-based acid tolerance system consisting of the glutaminase GlsA, glutamate decarboxylases GadA and GadB, and the glutamate/gamma-aminobutyric acid (GABA) antiporter GadC (44) **(Fig. 2).** Nonetheless, our findings suggest that at pH 5.5, MT123 shifts metabolic flux towards glutamate, which is then consumed through some unknown pathway. Further, our data suggest that this formation of glutamate is important for cellular fitness under mildly acidic conditions. We therefore asked whether glutamate is still being decarboxylated to GABA (70) but via a yet-uncharacterized decarboxylase(s). Our metabolomic data do, indeed, show significantly increased intracellular GABA abundances at pH 5.5 relative to pH 7.5 under both growth conditions **(Fig. S2).** In addition to decarboxylation of glutamate, GABA could also be formed from putrescine degradation. However, our growth medium was not supplemented with putrescine as a nitrogen source.

Additionally, we observed decreased abundance of PuuE—the reversible enzyme that produces GABA in this pathway—during growth at pH 5.5 under aerobic conditions **(Fig. 4A),** even though GABA also increases under these same conditions. Thus, we propose that, under mildly acidic conditions, MT123 might be producing GABA through the decarboxylation of glutamate via a yet-undescribed decarboxylase

The proteomic and mutant fitness data provide further insight into this speculative glutamate decarboxylase system. In our proteomic data, an enzyme annotated as an aspartate 1-decarboxylase (PanD)(71) is increased in abundance at pH 5.5 (relative to 7.5) during aerobic growth. This protein was also detected under denitrifying conditions; however, its log2FC did not reach our threshold of significance. We suggest that this decarboxylase might have a broader substrate specificity, allowing it to decarboxylate either glutamate or aspartate. Substrate promiscuity has been reported previously in amino acid decarboxylases (72,73). However, there were no significant mutant fitness effects associated with the gene (*panD*) encoding this enzyme. Thus, we propose that MT123 might have enzymes with redundant glutamate decarboxylase activities. For example, there is a second amino acid decarboxylase encoded in the MT123 genome, arginine/ornithine decarboxylase (AdiA/CadA/SpeF homolog in **Fig. 2**), as well several other decarboxylases annotated as having non-amino acid substrates.

We propose a mechanism for maintenance of cytoplasmic pH under mildly acidic conditions involving the consumption of a proton during glutamate decarboxylation by the yet-unknown glutamate decarboxylase (70). While glutaminase—an enzyme not observed in the MT123 genome **(Fig. 2A)**—is more typically implicated in acid tolerance, several reports have suggested a role glutamate synthase as well (74,75). Additional protons may also be consumed during glutamine conversion to glutamate by the glutamate dehydrogenase. For example, at pH 5, the fermentative bacterium *Clostridium beijerinckii* up-regulates the expression of the glutamate synthase operon(75). A remaining question is the identity of the antiporter system that is needed to exchange GABA for glutamate (76). However, prior work has suggested that endogenous glutamate may contribute partially, or fully, to acid tolerance in certain bacteria (77). For example, *Mycobacterium* species have glutamate-based acid tolerance systems, yet lack a GadC homolog (78). Decoupled intracellular GABA synthesis and efflux in response to low pH has also been reported in *Listeria monocytogenes* (79). In these organisms, it is speculated that GABA is recycled through the GABA shunt to succinate—involving the activities of a GABA aminotransferase and the succinate-semialdehyde dehydrogenase GabD/Sad. The MT123 genome encodes an in-tact GABA shunt (77). Interestingly, a GabD/Sad homolog (GFF664) increased in abundance at pH 5.5 under both growth conditions (aerobic: +4.9 log2FC, denitrifying: +3.2 Log2FC). However, the GABA aminotransferase of MT123, PuuE, is decreased in abundance—though was still detectable—under aerobic growth conditions **(Fig. 4A).** Some bacteria, like *Rhizobium leguminosarum*, have multiple transaminases with GABA transaminase activity (80). However, it is clear that more work is required to fully characterize these non-canonical acid tolerance systems in MT123 and these other bacteria.

There are two possible linkages between glutamate and BCAA metabolism that we considered. Perhaps the most obvious is the function of glutamate as an amino group donor to (3S)-3-methyl-2-oxopentanoate, 4-methyl-2-oxopentanoate, 3-methyl-2-oxobutanoate to yield isoleucine, leucine, and valine, respectively (81,82). Thus, BCAA biosynthesis consumes glutamate, a metabolite critical for MT123 acclimation to growth under acidic conditions. On the surface, this could explain the positive *Δfitness* values for several genes involved in BCAA biosynthesis **(Table 1).** However, this model does not explain (1) why exogenous isoleucine, leucine, and valine are toxic to MT123 under acidic conditions **(Fig. 8)** nor (2) the positive fitness values of genes encoding BCAA transporters in MT123 genomes alongside decreased abundances of these transport proteins **(Table 1).** Instead, we suggest that isoleucine, leucine, and valine accumulate intracellularly—through a yet-unknown mechanism—during growth under acidic conditions, an observation supported by our metabolomic data **(Fig. 5A-C).** Previously, diverse stress conditions have been shown to dysregulate BCAA biosynthesis in bacteria (83–85). As described earlier, leucine—and possibly also valine and isoleucine—are effectors of the global regulator Lrp (57). The MT123 genome encodes three Lrp homologs, which in other bacteria, regulate amino acid biosynthesis and degradation as well as other metabolic functions (86). In *E. coli*, the Lrp regulon includes both inducible and repressible genes, with leucine switching off or on these regulatory activities(87). With respect to leucine, Lrp is a repressible activator of the glutamate synthase gene (*glt*) in *E. coli* (87). However, in *E. coli* and other gram-negative bacteria, Lrp also appears to be responsive to other amino acids, including isoleucine, leucine, and valine (57,58,88). Thus, we suggest that the observed decreases in the abundances of Liv transporters and proteins involved in BCAA biosynthesis during growth at pH 5.5 may minimize this feedback inhibition on the glutamate biosynthesis required to sustain growth under acidic conditions.

Interestingly, more differentially abundant proteins were observed under aerobic conditions (432 proteins) compared to denitrifying conditions (158 proteins). This is despite the fact that we observed—both here and previously (24)—that MT123 has greater pH tolerance under aerobic conditions compared to denitrifying conditions. Protein synthesis (89) and turnover (90–92) are both ATP-dependent processes. However, under denitrifying conditions, growth at lower pH leads to accumulation of the protonophore nitrous acid, collapsing the cellular proton motive force (43). Prior studies have found that, in low quantities, nitrous acid can inhibit other ATP-dependent processes (42,93). Thus, we propose that the accumulation of nitrous acid during denitrifying growth at pH 5.5 contributes both to the reduced proteomic response and to the greater growth defects observed under denitrifying conditions.

In summary, we show how a multi-omics approach combined with pooled mutant fitness assays can be integrated to uncover novel insights into microbial acclimatation to environmental change in non-model systems. Prior studies on acid tolerance in gram-negative bacteria have been largely limited to human symbionts and pathogens (94), neglecting the diversity of tolerance mechanisms that may exist in free-living environmental bacteria that also experience wide environmental fluctuations in pH. Understanding these acid tolerance mechanisms in environmental bacteria is important for predicting how pH shifts can impact key microbially mediated environmental processes like nitrate removal from contaminated groundwater.

## DATA AVAILABILITY

The generated mass spectrometry proteomics data have been deposited to the ProteomeXchange Consortium via the PRIDE partner repository with the dataset identifier PXD064015 (95). Raw datafiles for the metabolomics experiments are available in MZML format through https://gnps2.org/status?task=29f23392a1414aa2a75002938e2fecec.

## Access details for REVIEWERS

Log in to the PRIDE website using the following details:

**Project accession:** PXD064015

**Token:** UmTQYuab9y66

Alternatively, reviewers can access the dataset by logging in to the PRIDE website using the following account details:

**Password:** bZSEhWUV9JO5

DIA-NN is freely available for download from https://github.com/vdemichev/DiaNN.

## ACKNOWLEDGEMENTS

This material by ENIGMA (Ecosystems and Networks Integrated with Genes and Molecular Assemblies) (http://enigma.lbl.gov), a Science Focus Area Program at Lawrence Berkeley National Laboratory, is based on work supported by the U.S. Department of Energy, Office of Science, Office of Biological and Environmental Research, under contract DE-AC02-05CH11231.

## REFERENCES

1. Majumdar D, Gupta N. Nitrate pollution of groundwater and associated human health disorders. Indian J Environ Health. 2000;42(1):28–39.

2. Shukla S, Saxena A. Sources and leaching of nitrate contamination in groundwater. Curr Sci [Internet]. 2020 [cited 2025 Feb 12];118(6):883–91. Available from: https://www.jstor.org/stable/27226382

3. Abascal E, Gómez-Coma L, Ortiz I, Ortiz A. Global diagnosis of nitrate pollution in groundwater and review of removal technologies. Sci Total Environ [Internet]. 2022 Mar 1 [cited 2025 Feb 12];810:152233. Available from: https://www.sciencedirect.com/science/article/pii/S0048969721073095

4. Zhao B, Sun Z, Liu Y. An overview of in-situ remediation for nitrate in groundwater. Sci Total Environ [Internet]. 2022 Jan 15 [cited 2025 Feb 12];804:149981. Available from: https://www.sciencedirect.com/science/article/pii/S0048969721050567

5. Calderer M, Gibert O, Martí V, Rovira M, de Pablo J, Jordana S, et al. Denitrification in presence of acetate and glucose for bioremediation of nitrate-contaminated groundwater. Environ Technol [Internet]. 2010 Jun 1 [cited 2025 Feb 12];31(7):799–814. Available from: 10.1080/09593331003667741

6. Moreels D, Crosson G, Garafola C, Monteleone D, Taghavi S, Fitts JP, et al. Microbial community dynamics in uranium contaminated subsurface sediments under biostimulated conditions with high nitrate and nickel pressure. Environ Sci Pollut Res [Internet]. 2008 Sep 1 [cited 2025 Feb 12];15(6):481–91. Available from: 10.1007/s11356-008-0034-z

7. Puigserver D, Herrero J, Carmona JM. Nitrate removal by combining chemical and biostimulation approaches using micro-zero valent iron and lactic acid. Sci Total Environ [Internet]. 2022 Oct 15 [cited 2025 Feb 12];843:156841. Available from: https://www.sciencedirect.com/science/article/pii/S0048969722039389

8. Safonov AV, Babich TL, Sokolova DS, Grouzdev DS, Tourova TP, Poltaraus AB, et al. Microbial Community and in situ Bioremediation of Groundwater by Nitrate Removal in the Zone of a Radioactive Waste Surface Repository. Front Microbiol [Internet]. 2018 Aug 23 [cited 2025 Feb 12];9. Available from: https://www.frontiersin.org/journals/microbiology/articles/10.3389/fmicb.2018.01985/full

9. Goff JL, Chen Y, Thorgersen MP, Hoang LT, Poole FL, Szink EG, et al. Mixed heavy metal stress induces global iron starvation response. ISME J. 2023;17(3):382–92.

10. Takem GE, Kuitcha D, Ako AA, Mafany GT, Takounjou-Fouepe A, Ndjama J, et al. Acidification of shallow groundwater in the unconfined sandy aquifer of the city of Douala, Cameroon, Western Africa: implications for groundwater quality and use. Environ Earth Sci [Internet]. 2015 Nov 1 [cited 2025 Mar 27];74(9):6831–46. Available from: 10.1007/s12665-015-4681-3

11. Appleyard S, Wong S, Willis-Jones B, Angeloni J, Watkins R. Groundwater acidification caused by urban development in Perth, Western Australia: source, distribution, and implications for management. Soil Res [Internet]. 2004 Sep 20 [cited 2025 Feb 12];42(6):579–85. Available from: https://www.publish.csiro.au/sr/SR03074

12. Kjøller C, Postma D, Larsen F. Groundwater Acidification and the Mobilization of Trace Metals in a Sandy Aquifer. Environ Sci Technol [Internet]. 2004 May 1 [cited 2025 Feb 12];38(10):2829–35. Available from: 10.1021/es030133v

13. Murdoch PS, Burns DA, Lawrence GB. Relation of Climate Change to the Acidification of Surface Waters by Nitrogen Deposition. Environ Sci Technol [Internet]. 1998 Jun 1 [cited 2025 Mar 27];32(11):1642–7. Available from: https://pubs.acs.org/doi/10.1021/es9708631

14. Murdoch PS, Stoddard JL. The role of nitrate in the acidification of streams in the Catskill Mountains of New York. Water Resour Res [Internet]. 1992 [cited 2025 Mar 27];28(10):2707–20. Available from: https://onlinelibrary.wiley.com/doi/abs/10.1029/92WR00953

15. Salminen JM, Petäjäjärvi SJ, Tuominen SM, Nystén TH. Ethanol-based *in situ* bioremediation of acidified, nitrate-contaminated groundwater. Water Res [Internet]. 2014 Oct 15 [cited 2025 Mar 27];63:306–15. Available from: https://www.sciencedirect.com/science/article/pii/S0043135414004424

16. Carreira C, Nunes RF, Mestre O, Moura I, Pauleta SR. The effect of pH on Marinobacter hydrocarbonoclasticus denitrification pathway and nitrous oxide reductase. JBIC J Biol Inorg Chem [Internet]. 2020 Oct 1 [cited 2025 Feb 14];25(7):927–40. Available from: 10.1007/s00775-020-01812-0

17. Liu B, Frostegård Å, Bakken LR. Impaired Reduction of N2O to N2 in Acid Soils Is Due to a Posttranscriptional Interference with the Expression of nosZ. mBio [Internet]. 2014 Jun 24 [cited 2025 Mar 27];5(3):10.1128/mbio.01383-14. Available from: https://journals.asm.org/doi/full/10.1128/mbio.01383-14

18. Olaya-Abril A, Hidalgo-Carrillo J, Luque-Almagro VM, Fuentes-Almagro C, Urbano FJ, Moreno-Vivián C, et al. Effect of pH on the denitrification proteome of the soil bacterium Paracoccus denitrificans PD1222. Sci Rep [Internet]. 2021 Aug 26 [cited 2025 Feb 14];11(1):17276. Available from: https://www.nature.com/articles/s41598-021-96559-2

19. Baumann B, van der Meer JR, Snozzi M, Zehnder AJ. Inhibition of denitrification activity but not of mRNA induction in Paracoccus denitrificans by nitrite at a suboptimal pH. Antonie Van Leeuwenhoek. 1997 Oct;72(3):183–9.

20. Du R, Peng Y, Cao S, Li B, Wang S, Niu M. Mechanisms and microbial structure of partial denitrification with high nitrite accumulation. Appl Microbiol Biotechnol [Internet]. 2016 Feb 1 [cited 2025 Feb 12];100(4):2011–21. Available from: 10.1007/s00253-015-7052-9

21. Pilegaard K. Processes regulating nitric oxide emissions from soils. Philos Trans R Soc B Biol Sci [Internet]. 2013 Jul 5 [cited 2024 Jul 5];368(1621):20130126. Available from: https://royalsocietypublishing.org/doi/abs/10.1098/rstb.2013.0126

22. Brooks SC. Waste characteristics of the former S-3 ponds and outline of uranium chemistry relevant to NABIR Field Research Center studies. NABIR Field Res Cent Oak Ridge Tenn. 2001;

23. Kämpfer P, Denger K, Cook AM, Lee ST, Jäckel U, Denner EBM, et al. Castellaniella gen. nov., to accommodate the phylogenetic lineage of Alcaligenes defragrans, and proposal of Castellaniella defragrans gen. nov., comb. nov. and Castellaniella denitrificans sp. nov. Int J Syst Evol Microbiol [Internet]. 2006 [cited 2025 Feb 14];56(4):815–9. Available from: https://www.microbiologyresearch.org/content/journal/ijsem/10.1099/ijs.0.63989-0

24. Goff JL, Szink EG, Durrence KL, Lui LM, Nielsen TN, Kuehl JV, et al. Genomic and environmental controls on Castellaniella biogeography in an anthropogenically disturbed subsurface. Environ Microbiome. 2024;19(1):26.

25. Spain AM, Peacock AD, Istok JD, Elshahed MS, Najar FZ, Roe BA, et al. Identification and Isolation of a Castellaniella Species Important during Biostimulation of an Acidic Nitrate- and Uranium-Contaminated Aquifer. Appl Environ Microbiol [Internet]. 2007 Aug [cited 2025 Feb 14];73(15):4892–904. Available from: https://journals.asm.org/doi/full/10.1128/aem.00331-07

26. Thorgersen MP, Ge X, Poole FL, Price MN, Arkin AP, Adams MWW. Nitrate-Utilizing Microorganisms Resistant to Multiple Metals from the Heavily Contaminated Oak Ridge Reservation. Appl Environ Microbiol [Internet]. 2019 Aug 14 [cited 2025 Feb 14];85(17):e00896–19. Available from: https://journals.asm.org/doi/full/10.1128/aem.00896-19

27. Widdel F, Bak F. Gram-Negative Mesophilic Sulfate-Reducing Bacteria. In: Balows A, Trüper HG, Dworkin M, Harder W, Schleifer KH, editors. The Prokaryotes: A Handbook on the Biology of Bacteria: Ecophysiology, Isolation, Identification, Applications [Internet]. New York, NY: Springer; 1992 [cited 2025 Feb 14]. p. 3352–78. Available from: 10.1007/978-1-4757-2191-1_21

28. McGinnis S, Madden TL. BLAST: at the core of a powerful and diverse set of sequence analysis tools. Nucleic Acids Res [Internet]. 2004 Jul 1 [cited 2025 Mar 4];32(suppl_2):W20–5. Available from: 10.1093/nar/gkh435

29. Wetmore KM, Price MN, Waters RJ, Lamson JS, He J, Hoover CA, et al. Rapid Quantification of Mutant Fitness in Diverse Bacteria by Sequencing Randomly Bar-Coded Transposons. mBio [Internet]. 2015 May 12 [cited 2025 Feb 14];6(3):10.1128/mbio.00306-15. Available from: https://journals.asm.org/doi/full/10.1128/mbio.00306-15

30. Thorgersen MP, Lancaster WA, Ge X, Zane GM, Wetmore KM, Vaccaro BJ, et al. Mechanisms of Chromium and Uranium Toxicity in Pseudomonas stutzeri RCH2 Grown under Anaerobic Nitrate-Reducing Conditions. Front Microbiol [Internet]. 2017 Aug 10 [cited 2025 Mar 28];8. Available from: https://www.frontiersin.org/journals/microbiology/articles/10.3389/fmicb.2017.01529/full

31. Chen Y, Gin J, Petzold CJ. Alkaline-SDS cell lysis of microbes with acetone protein precipitation for proteomic sample preparation in … 2023 Jan 18 [cited 2025 Jun 25]; Available from: https://www.protocols.io/view/alkaline-sds-cell-lysis-of-microbes-with-acetone-p-b2raqd2e

32. Yan Chen, Jennifer Gin, Christopher J Petzold. Discovery proteomic (DIA) LC-MS/MS data acquisition and analysis V.2. 2022.

33. Demichev V, Messner CB, Vernardis SI, Lilley KS, Ralser M. DIA-NN: neural networks and interference correction enable deep proteome coverage in high throughput. Nat Methods. 2020 Jan 1;17(1):41–4.

34. Ahrné E, Molzahn L, Glatter T, Schmidt A. Critical assessment of proteome-wide label-free absolute abundance estimation strategies. PROTEOMICS. 2013;13(17):2567–78.

35. Silva JC, Gorenstein MV, Li GZ, Vissers JPC, Geromanos SJ. Absolute quantification of proteins by LCMSE: a virtue of parallel MS acquisition. Mol Cell Proteomics MCP. 2006 Jan;5(1):144–56.

36. Sequeira JC, Rocha M, Alves MM, Salvador AF. UPIMAPI, reCOGnizer and KEGGCharter: Bioinformatics tools for functional annotation and visualization of (meta)-omics datasets. Comput Struct Biotechnol J. 2022 Jan 1;20:1798–810.

37. Yao Y, Sun T, Wang T, Ruebel O, Northen T, Bowen BP. Analysis of Metabolomics Datasets with High-Performance Computing and Metabolite Atlases. Metabolites [Internet]. 2015 Jul 20 [cited 2024 Jul 19];5(3):431–42. Available from: https://www.ncbi.nlm.nih.gov/pmc/articles/PMC4588804/

38. Mahler BJ, Bourgeais R. Dissolved oxygen fluctuations in karst spring flow and implications for endemic species: Barton Springs, Edwards aquifer, Texas, USA. J Hydrol. 2013 Nov 15;505:291–8.

39. Wenner E, Sanger D, Arendt M, Holland AF, Chen Y. Variability in Dissolved Oxygen and Other Water-Quality Variables Within the National Estuarine Research Reserve System. J Coast Res. 2004 Nov 1;(10045):17–38.

40. Mark B. Smith, Andrea M. Rocha, Chris S. Smillie, Scott W. Olesen, Charles Paradis, Liyou Wu, et al. Natural bacterial communities serve as quantitative geochemical biosensors. mBio. 2015;6(3):e00326–15.

41. Sijbesma WF, Almeida JS, Reis MA, Santos H. Uncoupling effect of nitrite during denitrification by Pseudomonas fluorescens: An in vivo 31P-NMR study. Biotechnol Bioeng. 1996;52(1):176–82.

42. Vadivelu VM, Yuan Z, Fux C, Keller J. The inhibitory effects of free nitrous acid on the energy generation and growth processes of an enriched nitrobacter culture. Environ Sci Technol. 2006 Jul 1;40(14):4442–8.

43. Zhou Y, Oehmen A, Lim M, Vadivelu V, Ng WJ. The role of nitrite and free nitrous acid (FNA) in wastewater treatment plants. Water Res. 2011 Oct 1;45(15):4672–82.

44. Mallick S, Das S. Acid-tolerant bacteria and prospects in industrial and environmental applications. Appl Microbiol Biotechnol. 2023 Jun 1;107(11):3355–74.

45. Xu Y, Zhao Z, Tong W, Ding Y, Liu B, Shi Y, et al. An acid-tolerance response system protecting exponentially growing Escherichia coli. Nat Commun. 2020 Mar 20;11(1):1496.

46. Mobley HLT. Urease. In: Mobley HL, Mendz GL, Hazell SL, editors. Helicobacter pylori: Physiology and Genetics [Internet]. Washington (DC): ASM Press; 2001 [cited 2025 Mar 4]. Available from: http://www.ncbi.nlm.nih.gov/books/NBK2417/

47. Goss TJ, Perez-Matos A, Bender RA. Roles of Glutamate Synthase, gltBD, and gltF in Nitrogen Metabolism of Escherichia coli and Klebsiella aerogenes. J Bacteriol. 2001 Nov 15;183(22):6607–19.

48. Pateman JA. Regulation of synthesis of glutamate dehydrogenase and glutamine synthetase in micro-organisms. Biochem J. 1969 Dec 1;115(4):769–75.

49. Fernie AR, Stitt M. On the Discordance of Metabolomics with Proteomics and Transcriptomics: Coping with Increasing Complexity in Logic, Chemistry, and Network Interactions Scientific Correspondence. Plant Physiol. 2012 Mar;158(3):1139–45.

50. Gupta M, Johnson ANT, Cruz ER, Costa EJ, Guest RL, Li SHJ, et al. Global protein turnover quantification in Escherichia coli reveals cytoplasmic recycling under nitrogen limitation. Nat Commun. 2024 Jul 13;15(1):5890.

51. Link H, Fuhrer T, Gerosa L, Zamboni N, Sauer U. Real-time metabolome profiling of the metabolic switch between starvation and growth. Nat Methods. 2015 Nov;12(11):1091–7.

52. Rahmanian M, Claus DR, Oxender DL. Multiplicity of Leucine Transport Systems in Escherichia coli K-12. J Bacteriol. 1973 Dec;116(3):1258–66.

53. Dutta S, Corsi ID, Bier N, Koehler TM. BrnQ-Type Branched-Chain Amino Acid Transporters Influence Bacillus anthracis Growth and Virulence. mBio. 2022 Feb 22;13(1):e0364021.

54. Stucky K, Hagting A, Klein JR, Matern H, Henrich B, Konings WN, et al. Cloning and characterization of brnQ, a gene encoding a low-affinity, branched-chain amino acid carrier in Lactobacillus delbrdückii subsp. lactic DSM7290. Mol Gen Genet MGG. 1995 Nov 1;249(6):682–90.

55. Belitsky BR. Role of Branched-Chain Amino Acid Transport in Bacillus subtilis CodY Activity. J Bacteriol. 2015 Mar 24;197(8):1330–8.

56. Kaiser JC, Heinrichs DE. Branching Out: Alterations in Bacterial Physiology and Virulence Due to Branched-Chain Amino Acid Deprivation. mBio. 2018 Sep 4;9(5):e01188–18.

57. Hart BR, Blumenthal RM. Unexpected Coregulator Range for the Global Regulator Lrp of Escherichia coli and Proteus mirabilis. J Bacteriol. 2011 Feb 9;193(5):1054–64.

58. Boulette ML, Baynham PJ, Jorth PA, Kukavica-Ibrulj I, Longoria A, Barrera K, et al. Characterization of alanine catabolism in Pseudomonas aeruginosa and its importance for proliferation in vivo. J Bacteriol. 2009 Oct;191(20):6329–34.

59. Haney SA, Platko JV, Oxender DL, Calvo JM. Lrp, a leucine-responsive protein, regulates branched-chain amino acid transport genes in Escherichia coli. J Bacteriol. 1992 Jan;174(1):108–15.

60. Thompson MG, Blake-Hedges JM, Cruz-Morales P, Barajas JF, Curran SC, Eiben CB, et al. Massively parallel fitness profiling reveals multiple novel enzymes in *Pseudomonas putida* lysine metabolism. mBio. 2019;10(3):e02577–18.

61. Gelfand DH, Steinberg RA. Escherichia coli mutants deficient in the aspartate and aromatic amino acid aminotransferases. J Bacteriol. 1977 Apr;130(1):429–40.

62. Baker TI, Crawford IP. Anthranilate Synthetase: PARTIAL PURIFICATION AND SOME KINETIC STUDIES ON THE ENZYME FROM ESCHERICHIA COLI. J Biol Chem. 1966 Dec 10;241(23):5577–84.

63. Kozlowski MC, Tom NJ, Seto CT, Sefler AM, Bartlett PA. Chorismate-utilizing enzymes isochorismate synthase, anthranilate synthase, and p-aminobenzoate synthase: mechanistic insight through inhibitor design. J Am Chem Soc. 1995 Mar;117(8):2128–40.

64. Powell JT, Morrison JF. The Purification and Properties of the Aspartate Aminotransferase and Aromatic-Amino-Acid Aminotransferase from Escherichia coli. Eur J Biochem. 1978;87(2):391–400.

65. Pérez-Arellano I, Rubio V, Cervera J. Dissection of *Escherichia coli* glutamate 5-kinase: Functional impact of the deletion of the PUA domain. FEBS Lett. 2005 Dec 19;579(30):6903– 8.

66. Jiang P, Pioszak AA, Ninfa AJ. Structure−Function Analysis of Glutamine Synthetase Adenylyltransferase (ATase, EC 2.7.7.49) of Escherichia coli. Biochemistry. 2007 Apr 1;46(13):4117–32.

67. Bergmeyer HUi. Methods of Enzymatic Analysis V2. Elsevier; 2012. 640 p.

68. Niu H, Li T, Du Y, Lv Z, Cao Q, Zhang Y. Glutamate Transporters GltS, GltP and GltI Are Involved in Escherichia coli Tolerance In Vitro and Pathogenicity in Mouse Urinary Tract Infections. Microorganisms. 2023 Apr 29;11(5):1173.

69. Krastel K, Senadheera DB, Mair R, Downey JS, Goodman SD, Cvitkovitch DG. Characterization of a Glutamate Transporter Operon, glnQHMP, in Streptococcus mutans and Its Role in Acid Tolerance. J Bacteriol. 2010 Feb;192(4):984–93.

70. De Biase D, Pennacchietti E. Glutamate decarboxylase-dependent acid resistance in orally acquired bacteria: function, distribution and biomedical implications of the operon. Mol Microbiol. 2012;86(4):770–86.

71. Tang Y, Liu Y, Chen Y, Zhang W, Zhao J, He S, et al. A review: Research progress on microplastic pollutants in aquatic environments. Sci Total Environ. 2021 Apr 20;766:142572.

72. Richardson G, Ding H, Rocheleau T, Mayhew G, Reddy E, Han Q, et al. An examination of aspartate decarboxylase and glutamate decarboxylase activity in mosquitoes. Mol Biol Rep. 2010 Oct 1;37(7):3199–205.

73. Wang Y, Xu H, White RH. β-Alanine Biosynthesis in Methanocaldococcus jannaschii. J Bacteriol. 2014 Aug;196(15):2869–75.

74. Djoko KY, Phan MD, Peters KM, Walker MJ, Schembri MA, McEwan AG. Interplay between tolerance mechanisms to copper and acid stress in *Escherichia coli*. Proc Natl Acad Sci. 2017 Jun 27;114(26):6818–23.

75. Reeve BWP, Reid SJ. Glutamate and histidine improve both solvent yields and the acid tolerance response of Clostridium beijerinckii NCP 260. J Appl Microbiol. 2016 May 1;120(5):1271–81.

76. Ma D, Lu P, Yan C, Fan C, Yin P, Wang J, et al. Structure and mechanism of a glutamate–GABA antiporter. Nature. 2012 Mar;483(7391):632–6.

77. Feehily C, Karatzas K a. g. Role of glutamate metabolism in bacterial responses towards acid and other stresses. J Appl Microbiol. 2013;114(1):11–24.

78. Rai R, Mathew BJ, Chourasia R, Singh AK, Chaurasiya SK. Glutamate decarboxylase confers acid tolerance and enhances survival of mycobacteria within macrophages. J Biol Chem. 2025 Feb 21;301(4):108338.

79. Karatzas KAG, Brennan O, Heavin S, Morrissey J, O’Byrne CP. Intracellular Accumulation of High Levels of γ-Aminobutyrate by Listeria monocytogenes 10403S in Response to Low pH: Uncoupling of γ-Aminobutyrate Synthesis from Efflux in a Chemically Defined Medium. Appl Environ Microbiol. 2010 Jun;76(11):3529–37.

80. Prell J, Boesten B, Poole P, Priefer UB. The Rhizobium leguminosarum bv. viciae VF39 γ-aminobutyrate (GABA) aminotransferase gene (gabT) is induced by GABA and highly expressed in bacteroidsThe GenBank accession number for the sequence determined in this work is AF335502. Microbiology. 2002;148(2):615–23.

81. Kuramitsu S, Ogawa T, Ogawa H, Kagamiyama H. Branched-Chain Amino Acid Aminotransferase of Escherichia coli: Nucleotide Sequence of the ilvE Gene and the Deduced Amino Acid Sequence1. J Biochem (Tokyo). 1985 Apr 1;97(4):993–9.

82. Yamamoto K, Tsuchisaka A, Yukawa H. Branched-Chain Amino Acids. In: Yokota A, Ikeda M, editors. Amino Acid Fermentation [Internet]. Tokyo: Springer Japan; 2017 [cited 2025 May 4]. p. 103–28. Available from: 10.1007/10_2016_28

83. Grothe S, Krogsrud RL, McClellan DJ, Milner JL, Wood JM. Proline transport and osmotic stress response in Escherichia coli K-12. J Bacteriol. 1986 Apr;166(1):253–9.

84. Tuite NL, Fraser KR, O’Byrne CP. Homocysteine Toxicity in Escherichia coli Is Caused by a Perturbation of Branched-Chain Amino Acid Biosynthesis. J Bacteriol. 2005 Jul;187(13):4362–71.

85. Jozefczuk S, Klie S, Catchpole G, Szymanski J, Cuadros-Inostroza A, Steinhauser D, et al. Metabolomic and transcriptomic stress response of Escherichia coli. Mol Syst Biol. 2010 Jan;6(1):364.

86. Brinkman AB, Ettema TJG, De Vos WM, Van Der Oost J. The Lrp family of transcriptional regulators. Mol Microbiol. 2003;48(2):287–94.

87. Cho BK, Barrett CL, Knight EM, Park YS, Palsson BØ. Genome-scale reconstruction of the Lrp regulatory network in Escherichia coli. Proc Natl Acad Sci. 2008 Dec 9;105(49):19462–7.

88. McFarland KA, Dorman CJ. Autoregulated expression of the gene coding for the leucine-responsive protein, Lrp, a global regulator in Salmonella enterica serovar Typhimurium. Microbiol Read Engl. 2008 Jul;154(Pt 7):2008–16.

89. Lengyel P, Söll D. Mechanism of protein biosynthesis. Bacteriol Rev [Internet]. 1969 Jun [cited 2025 May 13];33(2):264–301. Available from: https://journals.asm.org/doi/10.1128/br.33.2.264-301.1969

90. Baker TA, Sauer RT. ATP-dependent proteases of bacteria: recognition logic and operating principles. Trends Biochem Sci. 2006 Dec 1;31(12):647–53.

91. Bittner LM, Arends J, Narberhaus F. Mini review: ATP-dependent proteases in bacteria. Biopolymers. 2016;105(8):505–17.

92. Lee C, Schwartz MP, Prakash S, Iwakura M, Matouschek A. ATP-Dependent Proteases Degrade Their Substrates by Processively Unraveling Them from the Degradation Signal. Mol Cell. 2001 Mar 1;7(3):627–37.

93. Huang H, Liao J, Zheng X, Chen Y, Ren H. Low-level free nitrous acid efficiently inhibits the conjugative transfer of antibiotic resistance by altering intracellular ions and disabling transfer apparatus. Water Res [Internet]. 2019 Jul 1 [cited 2025 May 13];158:383–91. Available from: https://www.sciencedirect.com/science/article/pii/S0043135419303604

94. Liu Y, Tang H, Lin Z, Xu P. Mechanisms of acid tolerance in bacteria and prospects in biotechnology and bioremediation. Biotechnol Adv. 2015 Nov 15;33(7):1484–92.

95. Perez-Riverol Y, Bai J, Bandla C, García-Seisdedos D, Hewapathirana S, Kamatchinathan S, et al. The PRIDE database resources in 2022: a hub for mass spectrometry-based proteomics evidences. Nucleic Acids Res. 2022 Jan 7;50(D1):D543–52.

