## Supporting Information for "Contrasting effects of glutamate and branched-chain amino acid metabolism on acid tolerance in a *Castellaniella* isolate from acidic groundwater"

**SUPPLEMENTAL METHODS**

**RBTnSeq Library Construction.** We conjugated wild-type MT123 recipient cells with the *Escherichia coli* donor WM3064 carrying the *mariner* barcoded transposon vector pHLL250 (31). After conjugation for 12 hours on R2A plates supplemented with DAP (300 uM), we selected mutants on R2A agar plates supplemented with 25 µg/mL kanamycin. TnSeq was then performed to map transposon insertion sites and to link these to the unique DNA barcodes. The individual mutants are mapped to the MT123 genome based on the annotations provided in **Table S1**. **Table S2** relates the locus tags used for this study to the current version of the MT123 genome. The MT123 RB-TnSeq pooled library contains 420,430 uniquely barcoded mutants.

**Proteomic Analysis.** Protein was extracted from cell pellets and tryptic peptides were prepared by following established proteomic sample preparation protocol (1). Briefly, cell pellets were resuspended in Qiagen P2 Lysis Buffer (Qiagen, Germany) to promote cell lysis. Proteins were precipitated with addition of 1 mM NaCl and 4 x vol acetone, followed by two additional washes with 80% acetone in water. The recovered protein pellet was homogenized by pipetting mixing with 100 mM ammonium bicarbonate in 20% methanol. Protein concentration was determined by the DC protein assay (BioRad, USA). Protein reduction was accomplished using 5 mM tris 2-(carboxyethyl)phosphine (TCEP) for 30 min at room temperature, and alkylation was performed with 10 mM iodoacetamide (IAM; final concentration) for 30 min at room temperature in the dark. Overnight digestion with trypsin was accomplished with a 1:50 trypsin:total protein ratio. The resulting peptide samples were analyzed on an Agilent 1290 UHPLC system coupled to a Thermo Scientific Orbitrap Exploris 480 mass spectrometer for discovery proteomics (2). Briefly, peptide samples were loaded onto an Ascentis® ES-C18 Column (Sigma–Aldrich, USA) and were eluted from the column by using a 10 minute gradient from 98% solvent A (0.1 % FA in H2O) and 2% solvent B (0.1% FA in ACN) to 65% solvent A and 35% solvent B. Eluting peptides were introduced to the mass spectrometer operating in positive-ion mode and were measured in data-independent acquisition (DIA) mode with a duty cycle of 3 survey scans from m/z 380 to m/z 985 and 45 Tandem mass spectrometry (MS2) scans with precursor isolation width of 13.5 m/z to cover the mass range. DIA raw data files were analyzed by an integrated software suite DIA-NN. The databases used in the DIA-NN search (library-free mode) is the protein FASTA sequences based on the MT123 genome annotation provided in **Table S1,** plus the protein sequences of common proteomic contaminants. DIA-NN determines mass tolerances automatically based on first pass analysis of the samples with automated determination of optimal mass accuracies. The retention time extraction window was determined individually for all MS runs analyzed via the automated optimization procedure implemented in DIA-NN. Protein inference was enabled, and the quantification strategy was set to Robust LC = High Accuracy. Output main DIA-NN reports were filtered with a global false discovery rate set at 0.01 (FDR <= 0.01) on both the precursor level and protein group level. The Top3 method, which is the average MS signal response of the three most intense tryptic peptides of each identified protein, was used to quantify proteins in the samples (3,4). A jupyter notebook written in Python executed label-free quantification (LFQ) data analysis on the DIA-NN peptide quantification report, and the details of the analysis were described in the established protocol (5).

**REFERENCES**

1.	Chen Y, Gin J, Petzold CJ. Alkaline-SDS cell lysis of microbes with acetone protein precipitation for proteomic sample preparation in ... 2023 Jan 18 [cited 2025 Jun 25]; Available from: https://www.protocols.io/view/alkaline-sds-cell-lysis-of-microbes-with-acetone-p-b2raqd2e

2.	Yan Chen, Jennifer Gin, Christopher J Petzold. Discovery proteomic (DIA) LC-MS/MS data acquisition and analysis V.2. 2022.

3.	Silva JC, Gorenstein MV, Li GZ, Vissers JPC, Geromanos SJ. Absolute quantification of proteins by LCMSE: a virtue of parallel MS acquisition. Mol Cell Proteomics MCP. 2006 Jan;5(1):144–56.

4.	Ahrné E, Molzahn L, Glatter T, Schmidt A. Critical assessment of proteome-wide label-free absolute abundance estimation strategies. PROTEOMICS. 2013;13(17):2567–78.

5.	Chen Y, Petzold CJ. Label-free quantification (LFQ) proteomic data analysis from DIA-NN output files. 2022 Aug 2 [cited 2025 Jun 25]; Available from: https://www.protocols.io/view/label-free-quantification-lfq-proteomic-data-analy-b6f8rbrw

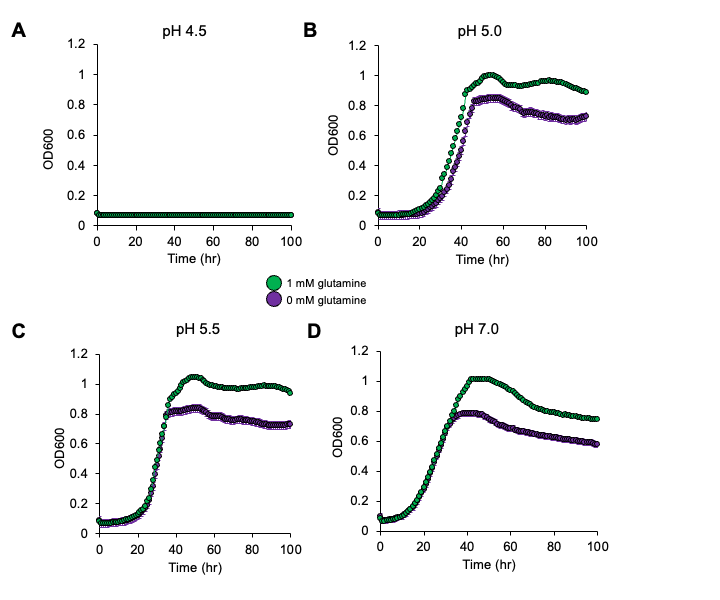

**Figure S1.** Growth of MT123 at (A) pH 5.0, (B) pH 5.5, (C) pH 6.0, (D) pH 7.0 with the addition of 1 mM glutamine under aerobic growth conditions. Individual points represent the average of three replicates. Error bars are present, though not always visible, and represent ±SD. The on-figure legend applies to all four panels.

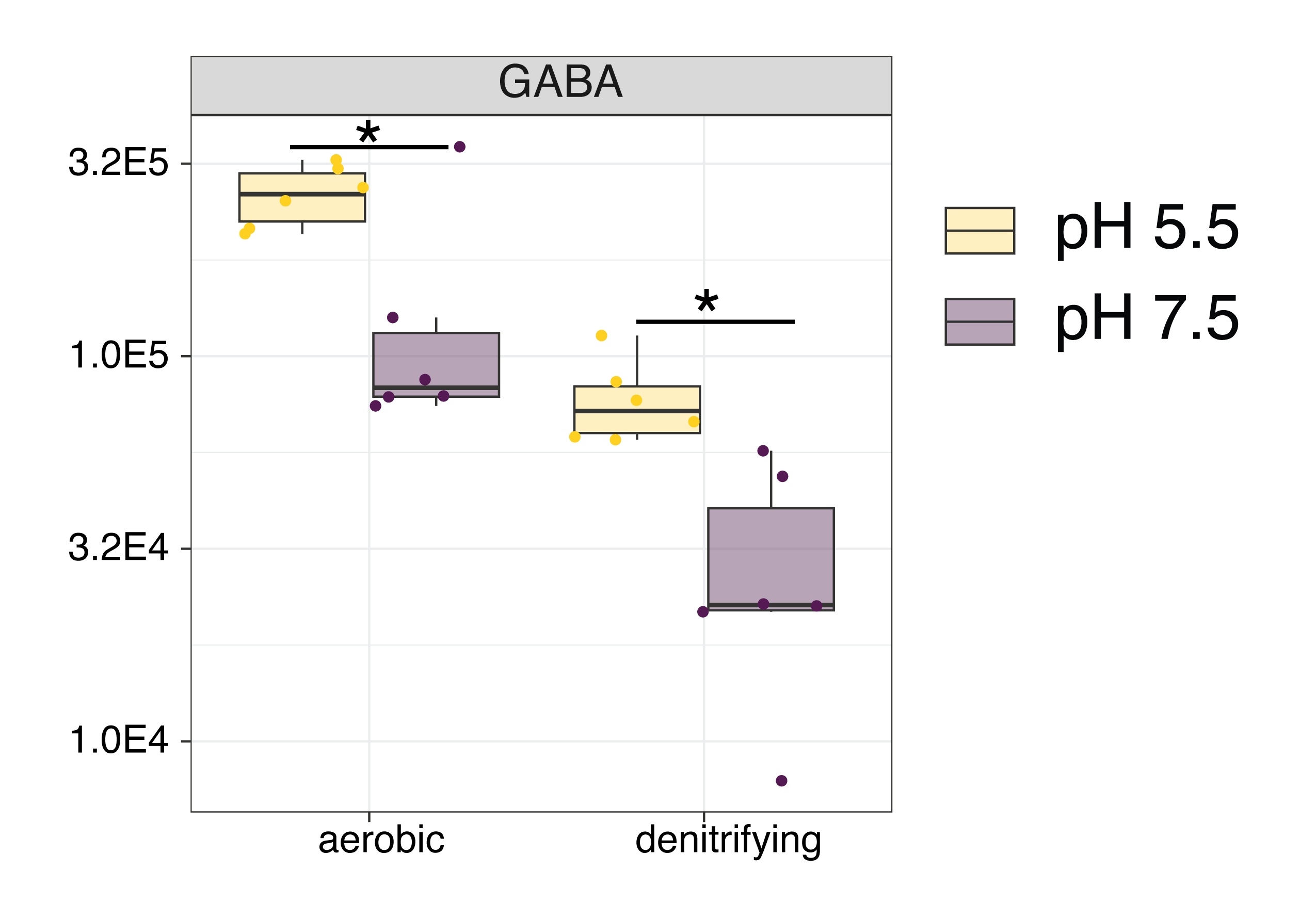

**Figure S2.** Cellular concentrations of 4-aminobutyric acid (GABA). Individual data points represent replicate experiments (n = 6). Statistical comparisons were performed with a Welch’s t-test using the Benjamini-Hochberg procedure to control for the false discovery rate (FDR). Asterisks (*) indicate an adjusted p-values <0.05.
